# Mathematical modeling unveils the timeline of CAR-T cell therapy and macrophage-mediated cytokine release syndrome

**DOI:** 10.1101/2024.04.23.590738

**Authors:** Daniela S. Santurio, Luciana R. C. Barros, Ingmar Glauche, Artur C. Fassoni

## Abstract

Chimeric antigen receptor (CAR)-T cell therapy holds significant potential for cancer treatment, although disease relapse and cytokine release syndrome (CRS) remain as frequent clinical challenges. To better understand the mechanisms underlying the temporal dynamics of CAR-T cell therapy response and CRS, we developed a novel multi-layer mathematical model incorporating antigen-mediated CAR-T cell expansion, antigen-negative resistance, and macrophage-associated cytokine release. Three key mechanisms of macrophage activation are considered: release of damage-associated molecular patterns, antigen-binding mediated activation, and CD40-CD40L contact. The model accurately describes 25 patient time courses with different responses and IL-6 cytokine kinetics. We successfully link the dynamic shape of the response to interpretable model parameters and investigate the influence of CAR-T cell dose and initial tumor burden on the occurrence of CRS and treatment outcome. By disentangling the timeline of macrophage activation, the model identified distinct contributions of each activation mechanism, suggesting the CD40-CD40L axis as a major driver of CRS and a clinically feasible target to control the activation process and modulate cytokine peak height. Our multi-layer model provides a comprehensive framework for understanding the complex interactions between CAR-T cells, tumor cells, and macrophages during therapy.

## Introduction

Immunotherapy with chimeric antigen receptor T (CAR-T) cells is a groundbreaking approach that harnesses the power of the immune system to fight hematological and non-hematological malignancies [1]. Genetic engineering enables T cells to express chimeric antigen receptors (CARs), enhancing their ability to specifically recognize and eliminate cancer cells. This remarkable efficacy in treating certain types of blood cancers made CAR-T cell therapy a promising avenue in oncology, with new CAR designs entering clinical trials for several indications, including solid tumors [2].

The short-term response to CAR-T cell therapy is characterized by typical multiphasic dynamics, with the initial distribution phase being marked by a decline of injected cells due to their death and migration to tumor site [3, 4, 5]. The subsequent expansion phase encompasses tumor-killing and a massive proliferation of CAR-T cells mediated by antigen recognition. In the following contraction phase, exhausted CAR-T cells are eliminated, and a fraction of long-lived memory CAR-T cells remain in the final persistence phase. Using mathematical models we and others showed how this multiphasic dynamics emerge from the interaction of different CAR-T cell phenotypes with tumor cells [3, 4, 5].

While CAR-T cell therapy is as a paradigm-shifting modality for cancer treatment, it faces substantial challenges, with 30-60% of patients experiencing recurrence within one year post-treatment [6]. Its effectiveness is linked to biological mechanisms of CAR-T cells, such as expansion, cytotoxicity, and memory formation, all of which are influenced by antigen recognition. Consequently, the density of antigens presented by tumor cells constitutes a pivotal factor for therapy resistance and transient antigen loss under treatment pressure or permanent antigen loss due to mutations account for typical resistance mechanisms [7].

Besides resistance, major obstacles for CAR-T cell therapy are side effects, including cytokine release syndrome (CRS) and immune effector cell-associated neurotoxicity syndrome (ICANS) [8]. Affecting 13%-26% of patients, severe CRS is the most prevalent adverse event, with symptoms typically emerging 1-7 days post-infusion, commonly before the CAR-T cell peak [9, 10]. CRS is characterized by fever, hypotension, and respiratory insufficiency and is associated with rising levels of cytokines, including IL-6, IL-10, IL-8, interferon-*γ* (IFN-*γ*) and granulocyte–macrophage colony-stimulating factor (GM-CSF) [11, 12]. In many cases CRS severity is tied to tumor burden and sometimes to CAR-T cell dose [13]. CRS also influences patient survival, and severe CRS (grade 3-5) correlated with lower survival in multiple myeloma (MM) and acute lymphoid leukemia (ALL) patients [11, 12].

Preclinical studies suggest that CRS results from complex interactions between CAR-T and host cells, with macrophages as a primary source of released cytokines [14, 15, 16, 8]. Three mechanisms have been identified as major drivers of macrophage activation underlying CRS [8]. First, upon antigenbinding-mediated activation, CAR-T cells release inflammatory signals such as IFN-*γ* and GM-CSF that induce macrophage activation in a contact-independent manner. Second, following tumor cell targeting by CAR-T cells, damage-associated molecular patterns (DAMPs) released by pyroptotic tumor cells enhance macrophage activation through pattern-recognition receptors. Third, the interaction of CD40 expressed on activated macrophages with CD40L expressed on CAR-T cells further promotes macrophage activation in a contact-dependent manner.

Different aspects of CAR-T cell therapy were investigated by modeling approaches [17, 18, 19, 20] with a few focusing on CRS [21, 22, 23, 24]. However, the critical role of macrophages on cytokine release was rarely addressed [25]. It is currently unclear how the different macrophage-associated mechanisms influences the extent and the temporal dynamics of CRS, although these aspects are essential for clinical implementation of countermeasures. With refined models, therapeutic interventions can be tested, as for example modulating CAR-T cell dose or changes in preconditioning to reduce the initial tumor burden. The models can also describe supportive administration of different corticosteroids or antibodies to inhibit macrophage activation and cytokine release.

Here, we develop a multi-layer model of CAR-T cell therapy encompassing antigen-negative resistance and macrophage-associated cytokine release. The model consistently describes time courses of 25 patients with different malignancies and tumor responses. Explicitly including three mechanisms of macrophage activation (DAMPs release, antigen-binding, and CD40 contact), the model also fits to cytokine time courses of 15 patients. We dissect the different phases of macrophage activation, showing that each mechanism occurs at a different time point during therapy response, with different contributions. Our results provide insights on macrophage-mediated cytokine release and elicit testable hypotheses on therapeutic interventions to mitigate CRS in CAR-T cell therapy.

## Methods

### Mathematical model

#### Modeling CAR-T cell dynamics and tumor response

Starting from our previous model [5], we developed a simplified version aimed at describing, with a minimal number of parameters, the multiphasic dynamics of CAR-T cells as a process emerging from the interaction between CAR-T phenotypes and their binding to antigen-expressing tumor cells (Figure 1). The model consists of four compartments representing three CAR-T cell phenotypes, namely injected, expanders, and persisters, represented as *C*_*D*_(*t*), *C*_*E*_(*t*), and *C*_*P*_ (*t*), respectively, and a population of antigen-positive tumor cells, represented as *T*_*P*_ (*t*). Incorporating established concepts of T cell biology, we further assume that T cell response rates are based on their phenotype - injected cells become active, activated cells expand, and persistent memory cells are re-activated - and proportional to their binding to antigen-expressing target cells. The basic model is formally given by the following system of ordinary differential equations (ODEs):

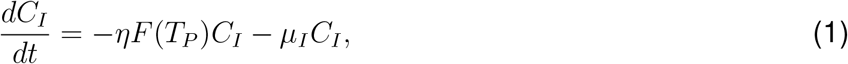

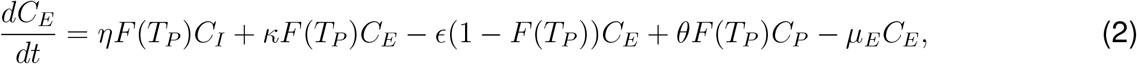

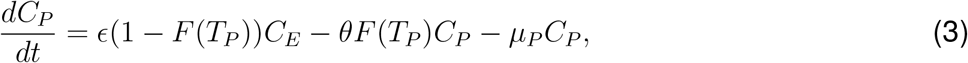

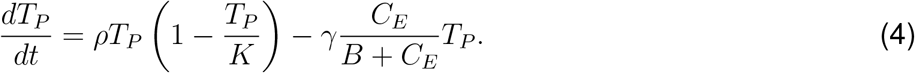

**Figure 1:**
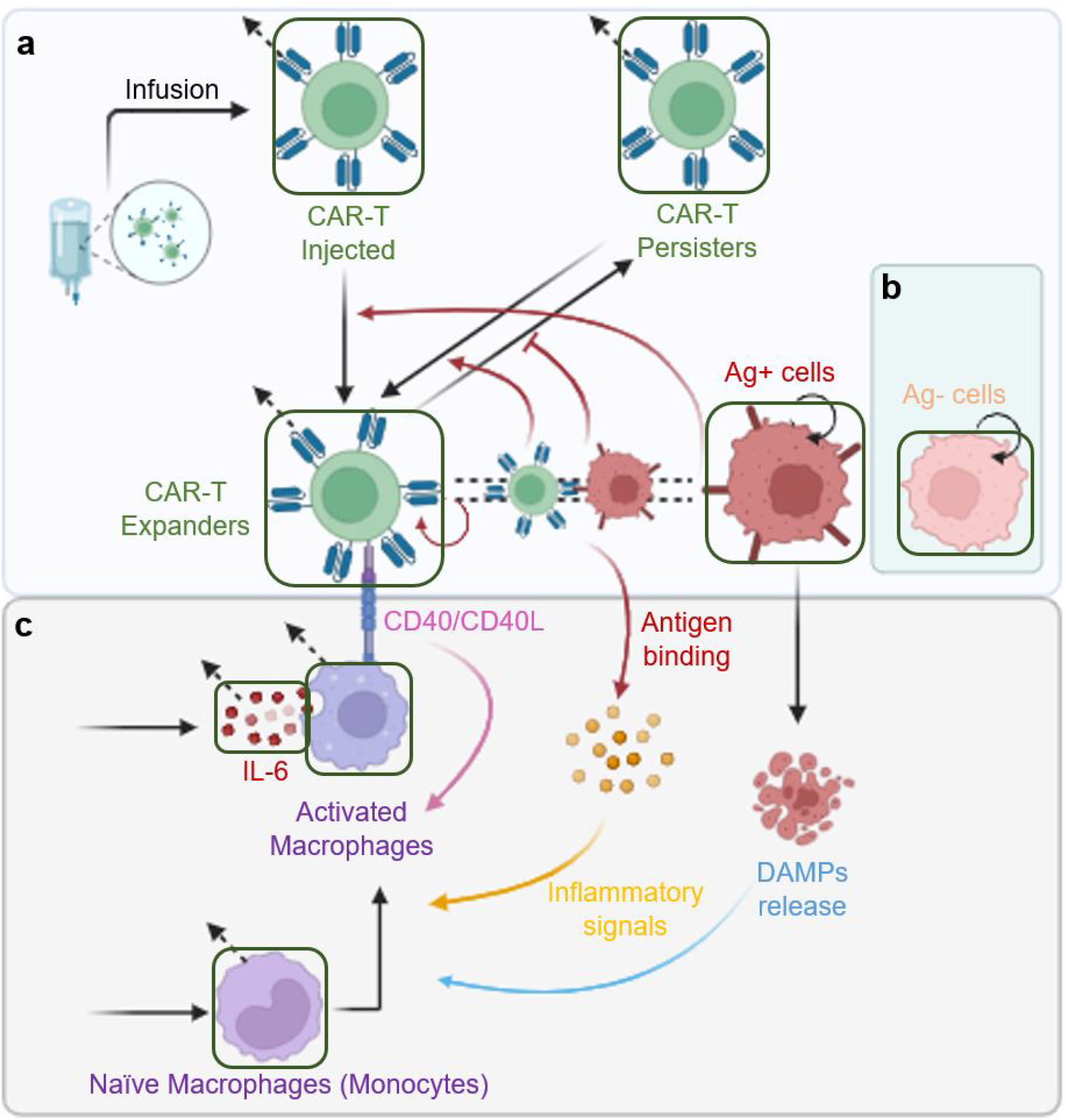
Model schematic description. **a** Model for CAR-T cell therapy considering injected (*C*_*I*_), expander (*C*_*E*_), and persister (*C*_*P*_) CAR-T cells, and antigen-positive (Ag+) tumor cells (*T*_*P*_). The phenotypic differentiation of CAR-T cells is driven by antigen recognition on the surface of tumor cells (red arrows). **b** The model is extended to describe patients who presented antigen-negative tumor relapses by including a compartment of antigen-negative (Ag-) tumor cells (*T*_*N*_). **c** The same model can be extended to describe macrophage-mediated cytokine release considering three different activation mechanisms: antigen-binding (upon antigen-recognition, activated CAR-T cells release inflammatory cytokines and molecules that activate naive macrophages, yellow arrow); DAMPs-release (CAR-T-mediated killing of the tumor leads to the release of damage-associated molecular patterns (DAMPs) that promote monocyte activation, blue arrow); CD40-contact (in a contact-dependent manner, activated macrophages expressing CD40 bind to CD40 ligand expressed by CAR-T cells and promote further macrophage activation (pink arrow).

The function *F* (*T*_*P*_) represents the antigen-receptor biding interaction and is described by a Michaelis-Menten functional response

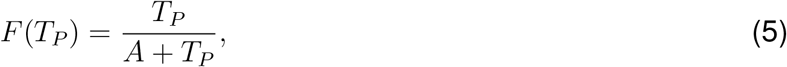

where *A* is the half-saturation constant. Following the approach in [26], we incorporated this function into all transition terms for CAR-T cells. This implies that the *per capita* rates of activation and proliferation of CAR-T expander cells and the activation rate of CAR-T persister cells are proportional to the antigen-binding function *F* (*T*_*P*_), while the rate of memory formation is proportional to 1 − *F* (*T*_*P*_). The model further assumes decay rates for each CAR-T cell phenotype, which include natural death and distribution throughout the body for injected cells (*µ*_*I*_), natural death and exhaustion for expanders (*µ*_*E*_), a slower natural death rate for persisters (*µ*_*P*_), and logistic growth for tumor cells with rate *ρ* and carrying capacity *K*. Following [27], the killing of tumor cells is described with a Holling type-II response with saturation for expanders cells; which also corresponds to simplifying our previous model with *ϑ* = 0 [5]. These Michaelis-Menten functional responses effectively capture the saturating nature of the antigen-receptor binding process, either by the limited number of antigen density in tumor cells (higher values of *A*) or by receptor density in CAR-T cells (higher values of *B*). An overview of model parameters is presented in Supplementary Table 1. Detailed information on model setup and parameter estimation is provided in Supplementary Texts S1-S3.

#### Modeling antigen-negative tumor relapse

The basic model is extended to include a compartment of antigen-negative tumor cells. Emerging from a simplification of a recent model for CAR-T cell therapy resistance [7], this second module has two ODEs describing the dynamics of antigen-positive (*T*_*P*_) and antigen-negative (*T*_*N*_) tumor cells, given by

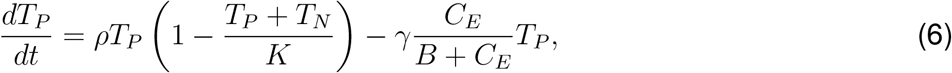

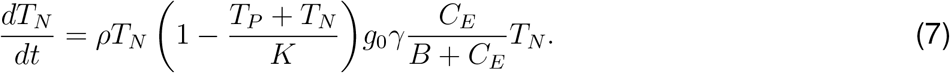

The killing of antigen-negative cells is multiplied by a fraction *g*_0_, which represents a reduced cytotoxicity due to antigen absence. A small but non-zero value for *g*_0_ is assumed to account for a minimal cytotoxic effect of CAR-T cells due to the bystander effect and/or the CAR-T cell’s endogenous TCR [7, 28, 29]. We also assume that antigen-negative tumor cells do not contribute to the antigen-dependent process of CAR-T cell activation, expansion, and memory recruitment. Therefore, the model for CAR-T cell therapy in a context with antigen-positive and antigen-negative tumor cells is formed by equations (1-3), (5) and (6-7).

#### Modeling macrophage-mediated cytokine release

Finally, a third layer is added to the model to describe cytokine dynamics and CRS. In line with widely accepted biology [15, 16, 30], we assume that cytokine release is mediated by macrophage activation. This activation and subsequent macrophage-recruitment is mainly driven by three mechanisms: i) upon target recognition, activated CAR-T cells release cytokines and soluble inflammatory mediators that activate macrophages, such as interferon-*γ*, granulocyte–macrophage colony-stimulating factor and tumor necrosis factor; ii) the killing of tumor cells releases DAMPs that further amplify macrophage activation; and iii) macrophage activation is also promoted by a contact-dependent mechanism, through the expression of CD40 and CD40-ligand by macrophages and CART-cells, respectively. We then assume that activated macrophages secrete inflammatory cytokines, including IL-6, chosen due to its significant role in CRS and the availability of patient time courses containing IL-6 kinetics alongside CAR-T cell responses. Under these assumptions, the model for macrophage activation and cytokine release reads as

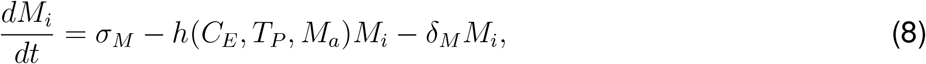

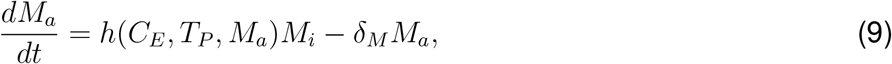

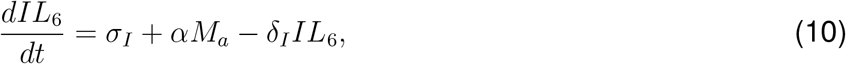

where *M*_*i*_ and *M*_*a*_ represent the naive (monocytes) and activated macrophages, respectively, and *IL*_6_ describes the IL-6 concentration. Furthermore, *σ*_*M*_ is the natural production rate of naive macrophages, *δ*_*M*_ is the death rate of macrophages, *σ*_*I*_ represents the endogenous production rate of IL-6, *δ*_*I*_ is the IL-6 natural decay rate, and *α* is the rate of IL-6 release by activated macrophages. The macrophage activation rate is given by

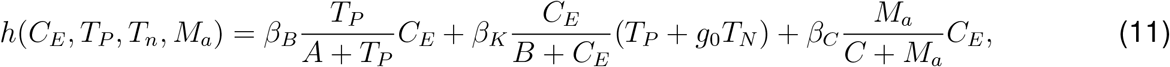

and encompasses the contribution of the three different mechanisms for macrophage activation: antigen-binding-mediated release of inflammatory signals by CAR-T cells (first term, proportional to CAR-T activation, with contribution *β*_*B*_), tumor-killing-mediated release of DAMPs (second term, proportional to tumor killing, with contribution *β*_*K*_), and CAR-T cell and macrophages contact (third term, Michaelis-Menten kinetics for CD40-CD40L binding, with contribution *β*_*C*_).

### Clinical data

We did an extensive search in the literature for datasets containing patient time courses with enough data points for fitting multiphasic CAR-T kinetics and IL 6 kinetics. We then included in our modeling all datasets satisfying these requirements. Individual pharmacokinetic profiles of patients undergoing anti-CD19 CAR-T cell therapy were digitized from [1, 31, 32, 33], and we collected the raw data disclosed in [34]. A detection limit was set for CAR-T at 25 copies/*µ*g DNA [34] or 2.5×10^5^ cells (for conversions see Supplementary Text S1) and relapse was defined as the absolute number of total tumor cells exceeding its initial tumor burden.

We incorporated data from [1] for two pediatric patients. Patient 1 received a total of 1.2×10^7^/kg CAR-T cells administered over three consecutive days, while Patient 2 received a single infusion of 1.4×10^6^/kg CAR-T cells. Although specific initial tumor burden data were not provided, CD19 expression analysis in bone marrow samples of Patient 2 revealed a pre-treatment blast population comprising approximately 7.7% CD19-cells and 92.3% CD19+ cells. In the absence of peripheral blood measurements, we assumed the same distribution of antigen-negative and positive cells to set the initial tumor burden. Patient 2 experienced a clinical relapse evident in the peripheral blood two months post-infusion, while Patient 1 achieved clinical remission that persisted for nine months.

The dataset from [31] includes a phase I study with pediatric B-ALL patients treated using a second-generation CAR with a 4-1BB intracellular co-stimulatory domain. Each patient received one CAR-T cell infusion, with initial tumor burden estimated from pre-therapy peripheral blood blast percentages. Progression-free survival (PFS) days was used as relapse times. The IL-6-profile was represented as a heatmap for the first month, and we computed the average value to capture the time series results of IL-6 concentrations in serum.

We used data from [32], patients 1, 2, 3, and 4 diagnosed with ALL received treatment using autologous CAR with CD28 intracellular co-stimulatory domain, while patients 6 and 7 were treated with autologous CAR featuring CD137 intracellular co-stimulatory domain. The time-profile of CD19+ cells in lymphocytes in the blood, measured by flow cytometry, revealed that within 2-8 months, patients experienced either progression or relapse of CD19+ leukemia cells.

Finally, we used data from [34], for adult chronic lymphocytic leukemia (CLL) patients treated with split-dose of autologous T-cells transduced with a CD19-directed CAR. With the exception of patient 22, who transformed to CD19-dim DLBCL, patients exhibited either complete or partial responses. For patients P01, P02 and P03, the initial CLL tumor burden and cytokine information (baseline and fold change) were extracted from [33] (respectively named UPN01, UPN03 and UPN02). Initial tumor burden on day -1 was available for UPN02 (P03), while for UPN01 (P01) and UPN03 (P02), only bone marrow data was provided, we assumed a similar approximation in blood due to its reasonable range.

An overview of patient and product characteristics and the corresponding infused doses and regimens are presented in Supplementary Table 2.

## Results

### The model accurately reproduces diverse patient-specific responses

The fundamental layer of our modeling approach is a minimal model describing CAR-T cell multiphasic dynamics and antigen-positive tumor responses (Figure 1a, equations (1-5)). This layer encompasses three CAR-T cell populations, namely injected, expander, and persister cells, and a population of antigen-positive tumor cells. The underlying model emerges as simplification from our previous model [5] with a reduction from 19 to 12 parameters, now integrating concepts of T-cell turnover [26] and plasticity [27, 35] which explain transitions between CAR-T cell phenotypes as functional responses to antigen-binding (Figure 1a).

Our modeling approach is based on the idea that the T-cell response is driven by activated, expanding cells that eliminate tumor cells, and persistent, long-lived memory cells that form a silent reservoir, which can be reactivated to maintain sustained remission [36]. Instead of fixed progenitor relationships between effector and memory T cells, our model assumes phenotypic plasticity between these cell types depending on the available antigen. Given that central memory and effector memory cells can also proliferate [37, 38], we group these phenotypes into a single population termed CAR-T expander cells, while long-term memory cells with low cytotoxicity are named CAR-T persister cells, consistent with the modeling framework proposed in [36]. The switch between these phenotypes depends on the antigen load. High antigen levels activate injected CAR-T cells, induce the proliferation of CAR-T expanders, and lead to conversion of persister cells into expander cells and inhibit the formation of persister cells.

Fitting the model (details provided in Supplementary Texts S1–S3) to n=19 time courses of CAR-T-treated patients with different malignancies (Figure 2, Supplementary Figures 1 and 2), shows that the model accurately describes both the CAR-T cell multiphasic dynamics as well as the different types of tumor responses.

**Figure 2:**
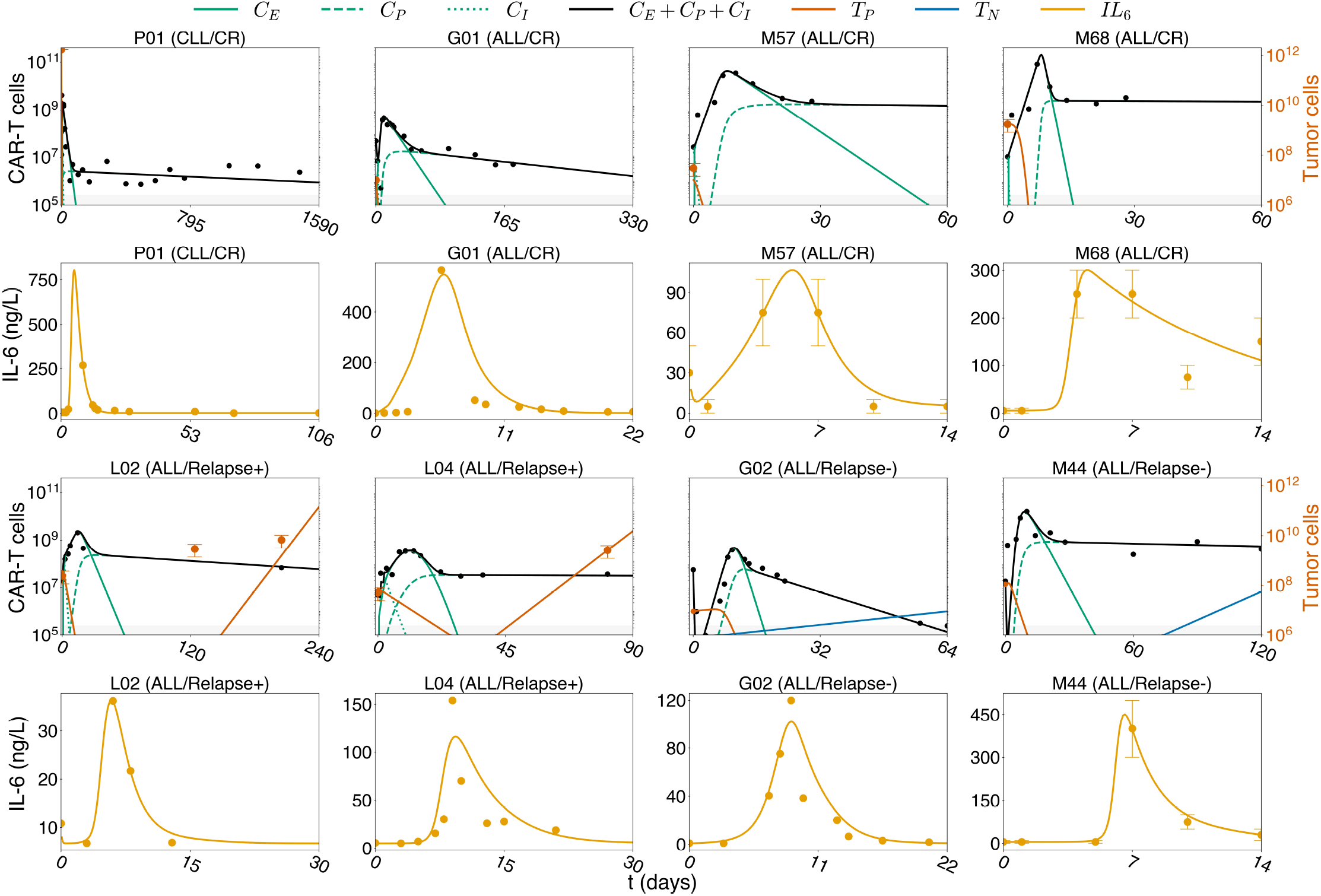
Model simulations. Model fits for selected patients that showed response to therapy (P01, G01, M57, and M68) or relapse either of antigen-positive tumor cells (L02, L04) or antigen-negative tumor cells (G02, and M44). See Supplementary Figures 1–4 for all 25 patients and Supplementary Texts S1–S3 for details on parameter estimation. Although patients M57 and M68 were simulated until their PFS day, plots show their dynamics for 60 days for clarity. Tumor cell error bars represent the range of WBCs used in scaling while error bars for IL-6 in patients M44, M57, and M68 encompass the range reported in [31]. The CAR-T cell detection threshold of 2.5 × 10^5^ cells is represented by the gray shaded area.

Tumor relapse occurs in 30-60% of patients within one year after CAR-T therapy [6] and many relapsed patients present antigen-negative tumor cells, which are more resistant to CAR-T cytotoxicity. To account for those cases, we add a second layer to the basic model, consisting of a population of antigen-negative tumor cells (Figure 1b, equation (7)). Due to the lack of antigen, we assume that these cells do not contribute to the antigen-mediated phenotypic changes in CAR-T cells. However, they are subject to a minimal cytotoxic effect by CAR-T cells due to the bystander effect and CAR-T cell endogenous T-cell receptor [28, 7]. The model was fitted to n=6 time courses of antigen-negative relapsed patients (Figure 2, Supplementary Figure 3). We confirm that the model quantitatively and qualitatively reflects the dynamics of antigen-negative relapses.

To detect early indications of severe CRS in CAR-T cell therapy, the concentrations of critical cytokines are usually monitored in parallel to treatment response, specifically IL-6. Typically, these cytokines follow a predictable kinetics: starting from a baseline concentration, they often peak before the CAR-T cells, and then return to baseline levels. It is increasingly evident that macrophages play a pivotal role in the release of cytokines in response to CAR-T cell therapy [14, 16, 8]. To quantitatively describe macrophage-mediated cytokine kinetics and gain a deeper understanding on the mechanisms underlying CRS, we add a third layer to our model including three compartments, namely naive macrophages, activated macrophages and IL-6 blood concentration (Figure 1c, equations (8-11)). We focus our attention on IL-6 as more time course data is available.

The main assumption of the model is that macrophage activation is driven by three different, well-described mechanisms: (i) the release of DAMPs by the killing of tumor cells, (ii) the release of inflammatory signals by CAR-T cells upon antigen recognition, and (iii) the contact of activated macrophages and CAR-T cells through the CD40-CD40L axis. In our model we further assume basal influx/production and death/decay rates for monocytes and cytokines, and that IL-6 is released by activated macrophages. We collected additional IL-6 time courses for a subset of 15 patients for which also CAR-T cell time courses were available. The model is fitted with excellent agreement between data and simulations (Figure 2, Supplementary Figure 4).

We conclude that our modular model structure accurately describes the main dynamical aspects of CAR-T cell therapy, explicitly accounting for CAR-T cell multiphasic dynamics, antigen-positive and negative relapses, and the occurrence of macrophage-mediated CRS.

### Different functional mechanisms shape the dynamics of therapy response

The response to CAR-T cell therapy unfolds over time, characterized by distinct phases in CAR-T cell dynamics. Mathematical approaches are suited to identify the functional mechanisms driving each re-sponse phase [5]. The fact that both CAR-T and tumor cell numbers change rapidly by different orders of magnitude after CAR-T cell infusion translates into mathematical arguments that allow simplifying the model equations for each phase and then identifying leading order effects. As detailed in Supplementary Text S4, each phase is approximated by an exponential function that is represented on the logarithmic scale as a straight line with characteristic slopes related to different model parameters and reflecting patient-specific values (Figure 3).

**Figure 3:**
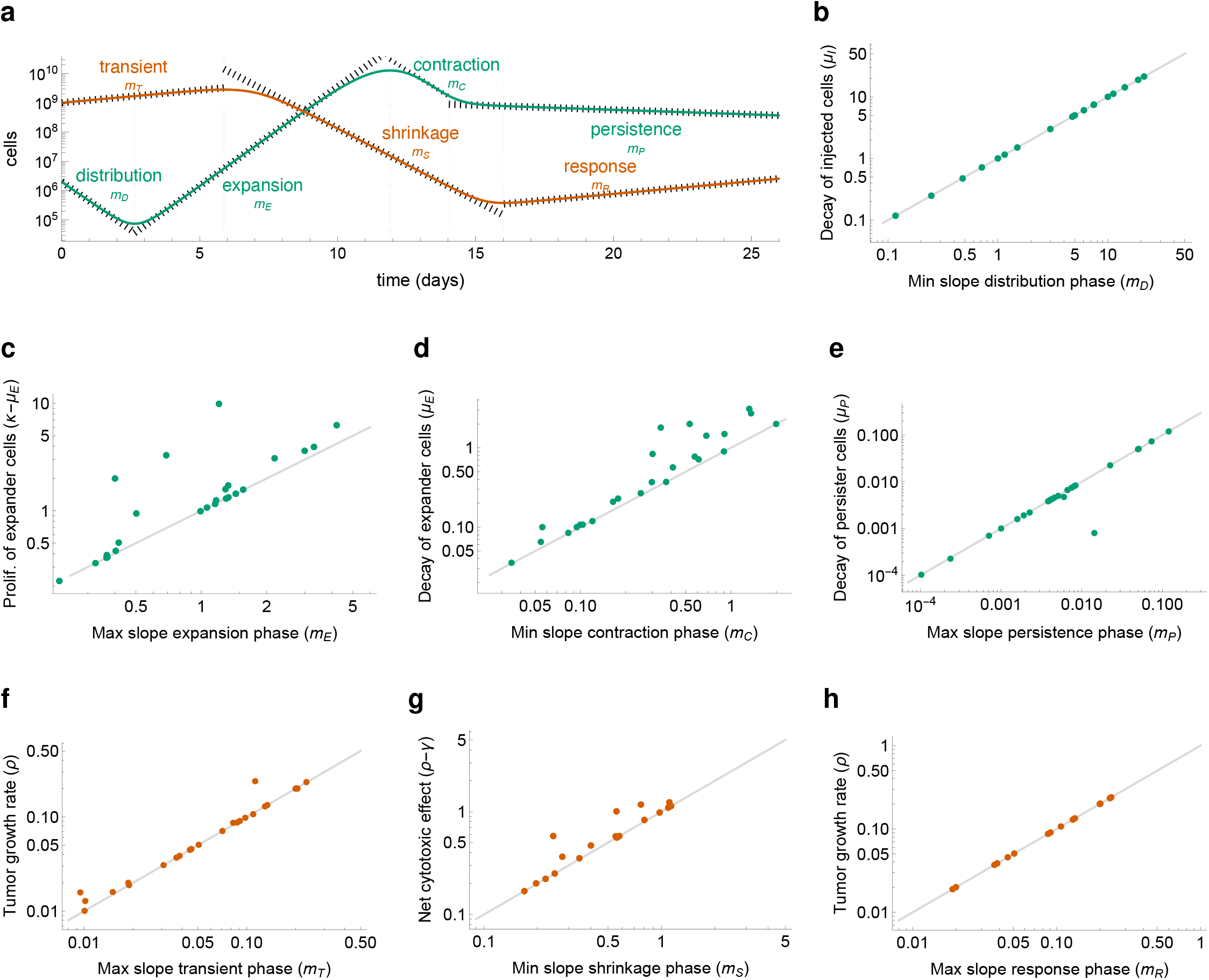
Identifying mechanisms driving therapy phases. **a** For each patient model fit, the distinct phases for CAR-T (green) and tumor (red) cell responses are identified and the characteristic slope is calculated and compared with the patient-specific mechanistic parameter driving the corresponding phase. **b-h** Comparison between the calculated slope and the mechanistic parameter for each phase and each patient. **b** Distribution phase: the calculated slope is the minimum (negative) slope of CAR-T cells, the mechanistic parameter is −*µ*_*I*_. **c** Expansion phase: the calculated slope is the maximum (positive) slope of CAR-T cells, the mechanistic parameter is *κ*−*µ*_*E*_. **d** Contraction phase: the calculated slope is the minimum (negative) slope of CAR-T cells, the mechanistic parameter is −*µ*_*E*_. **e** Persistence phase: the calculated slope is the maximum (negative) slope of CAR-T cells, the mechanistic parameter is −*µ*_*P*_. **f** Transient phase: the calculated slope is the maximum (positive) slope of tumor cells, the mechanistic parameter is *ρ*. **g** Shrinkage phase: the calculated slope is the minimum (negative) slope of tumor cells, the mechanistic parameter is *ρ* − *γ*. **h** Response phase: the calculated slope is the maximum (positive) slope of tumor cells for relapse patients only, the mechanistic parameter is *ρ*.

Comparing the slopes calculated from the model fits with the model parameter responsible for each exponential phase, our analysis shows that the four phases of CAR-T cell dynamics are driven by distinct mechanisms and cellular phenotypes (Figure 3a-e). The distribution phase is dominated by the decay of injected CAR-T cells, with the slope closely correlated to their decay rate −*µ*_*I*_. The subsequent expansion phase is driven by CAR-T expanders, with slope correlated to the net effect between their expansion and reduction due to death and exhaustion, represented as *κ* − *µ*_*E*_. The contraction phase is characterized by the exhaustion and death of CAR-T expander cells, with slope correlated to their depletion rate −*µ*_*E*_. The final persistence phase is dominated by the presence of residual persistent CAR-T cells, with slope closely correlated to their death rate −*µ*_*P*_.

We also identify three phases for tumor responses (Figure 3f-h). The first is a short and sometimes missing transient phase characterized by tumor growth while the injected CAR-T cells are being activated. Its slope is correlated to the tumor growth rate *ρ*. The second phase is the shrinkage phase, characterized by a strong decay in the tumor burden due to the CAR-T-mediated killing. Its slope is correlated to the difference between the tumor growth rate and CAR-T cytotoxic effect, *ρ* − *γ*. The last phase is the response phase, which may be either a re-expansion leading to relapse with slope closely correlated to the tumor growth rate *ρ*, or tumor extinction. The final outcome depends on an intricate balance of all pro- and anti-tumor parameters and whether or not the CAR-T cell-mediated killing reduced the tumor burden below a residual disease level.

The identification of model parameters driving the various phases of therapy response in clinical data not only enhances our mechanistic understanding but also proves indispensable for model fitting. As we demonstrate below, the model fitting is further complemented by identifying other parameters governing shape features of the model solutions.

In summary, we could successfully and uniquely link the different phases of the CAR-T and tumor cell response to the underlying mechanistic parameters of our mathematical model.

### Analysis of CAR-T and tumor cell dynamics enables informed parameter estimation

The preceding analysis enhances our understanding of the multiphasic dynamics following CAR-T cell administration and reveals a mapping between six characteristic slopes (*m*_*D*_, *m*_*E*_, *m*_*C*_, *m*_*P*_, *m*_*S*_, *m*_*P*_, see Figure 3) and six model parameters combinations (*µ*_*D*_, *κ* − *µ*_*E*_, *µ*_*E*_, *µ*_*P*_, *γ, ρ*), where each slope is approximated by the respective combination at first-order. In terms of parameter estimation, this allows for an informed initial guess for these parameters. However, such analysis does not elucidate how the remaining four characteristic shape features of therapy response, such as the CAR-T cell minimum after distribution, the CAR-T cell peak concentration (*C*_*max*_), the CAR-T cell persistence level, and the minimum tumor burden achieved by therapy (*T*_*min*_), correlate with model parameters, specially with the four remaining patient-specific parameters *A, B, η, ϵ*. To address this question, we conducted a qualitative sensitivity analysis. Technically, we assess the effect of each model parameter, one at a time, on these response hallmarks (Figure 4, Supplementary Figure 5).

**Figure 4:**
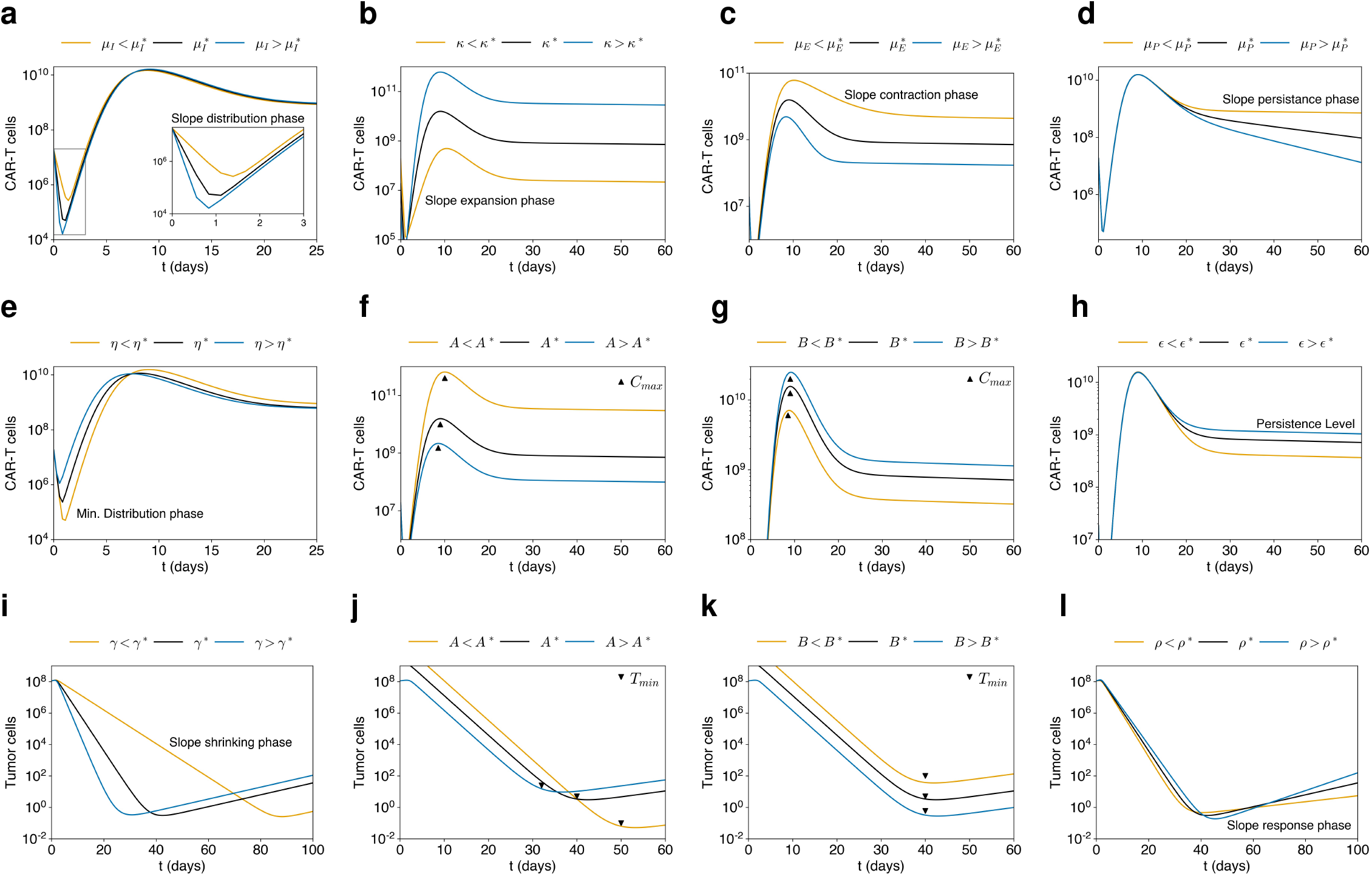
Mapping of mechanistic parameters on response characteristics. A qualitative sensitivity analysis identified a nonlinear mapping between ten different shape features of the model response (slopes *m*_*D*_, *m*_*E*_, *m*_*C*_, *m*_*P*_, *m*_*S*_, *m*_*P*_, CAR-T cell minimum, CAR-T cell peak concentration, minimum tumor load, and persistence level) to ten mechanistic parameters. This is illustrated by simulations for patient M44 (black) and alternative scenarios (blue and yellow) where one parameter is changed at a time. **a-d** The slopes in the distribution, expansion, and contraction phases are determined by the decay of injected CAR-T cells (*µ*_*I*_), the net expansion rate of CAR-T expander cells (*κ* − *µ*_*E*_), their death and exhaustion (*µ*_*E*_), and the decay of CAR-T persister cells (*µ*_*P*_). **e** The minimum level of CAR-T cells observed at the end of the distribution phase is determined by the activation rate of injected CAR-T cells (*η*). **f**,**g** The CAR-T cell peak depends on the saturation constants for the antigen binding (*A*) and tumor killing (*B*) functions. **h** The persistence level is mainly determined by the memory pool formation rate *ϵ*. **i**,**j** The slopes in the shrinkage and response phases of tumor dynamics are determined by CAR-T cell cytotoxicity (*γ*) and tumor growth rate (*ρ*), respectively. **k**,**l** The minimum tumor load is negatively correlated with the CAR-T cell peak and therefore is also determined by *A* and *B*. Although most of the response characteristics appear to have a first-order dependence on only one mechanistic parameter, this is not the case for the CAR-T cell peak and the minimum tumor burden, which also strongly depend on *κ, µ*_*E*_, *γ* and *ρ*. See also Supplementary Figure 5.

First, we found that the CAR-T cell minimum after the distribution phase is intrinsically related to the engraftment rate *η*, allowing us to obtain a first estimate for this parameter from the duration of the distribution phase, since the death rate of injected CAR-T cells is approximated by the distribution slope. We also identified that *C*_*max*_ and *T*_*min*_ depend on the expansion rate *κ*, the tumor growth rate *ρ*, the cytotoxic kill rate of CAR-T cells on tumor cells *γ*, the depletion rate *µ*_*E*_, and the half-saturation constants *A* and *B*, with higher *C*_*max*_ associated with lower *T*_*min*_. Considering that *κ, ρ, γ* and *µ*_*E*_ can be initially mapped by the characteristic slopes, this implies that initial estimates for *A* and *B* can be obtained from *C*_*max*_ and *T*_*min*_. From the biological point of view, this indicates that CAR-T cell expansion and tumor response are related to both product and patient-related characteristics, including the densities of antigens in tumor cells (*A*) and receptors in CAR-T cells (*B*). Finally, we observed that the CAR-T cell persistence level is closely linked to the memory formation rate *ϵ* and on the CAR-T cell peak itself (along with the parameters governing *C*_*max*_). Again, considering that the previous parameters were already mapped, this allows the estimation of *ϵ*.

This analysis then identifies a mapping between the ten different shape features of the model response (six characteristic slopes, the minimum tumor load, CAR-T cell minimum, peak concentration and persistence level) to ten model parameters (*µ*_*D*_, *κ, µ*_*E*_, *µ*_*P*_, *γ, ρ, η, ϵ, A* and *B*), and provides a rationale for estimating these parameters. The remaining two parameters *K* (tumor cell carrying capacity) and *θ* (persister cell recruitment rate) do not have major influence on the shape of the model solutions and therefore can be set to fixed, universal values. Based on these findings, we formulated a fitting strategy (Supplementary Text S2) that utilizes the clinical time courses to derive initial estimates for these 10 parameters and then employs an optimization algorithm to obtain optimal model fits for each patient (Figure 2). The efficacy of this strategy underscores the intimate relationship between the shape of the model solutions and the model parameters. Thus, our model can be regarded as a minimal model for delineating multiphasic responses, and the one-to-one mapping of data to mechanistic parameters ensures parameter identifiability in our fitting strategy.

In conclusion, we demonstrated how distinct model parameters dictate the shape of CAR-T and tumor cell response, providing a systematic approach for parameter estimation and revealing minimal redundancy in the model parameters.

### Clinical data suggests poor predictive value of patient-specific factors on IL-6 fold change, but strong temporal connection

So far we illustrated how each feature of the dynamical therapy response corresponds to distinct functional parameters of our model. However, it is well established that certain pharmacological and clinical factors such as CAR-T cell dose, CAR-T cell peak and its time, and initial tumor burden are associated with therapy response or can serve as indicators of severe side-effects [39, 40, 41]. Our modeling approach allows to systematically correlate these quantitative estimates with predicted clinical outcomes. Therefore, for each patient’s time course, we evaluated and correlated key parameters, including the fold change in IL-6 concentration from baseline to peak, the fold change for CAR-T cell counts from dose to peak, peak times for CAR-T cells, IL-6, and macrophages, as well as the tumor burden at days 28 and 90 (Figure 5).

**Figure 5:**
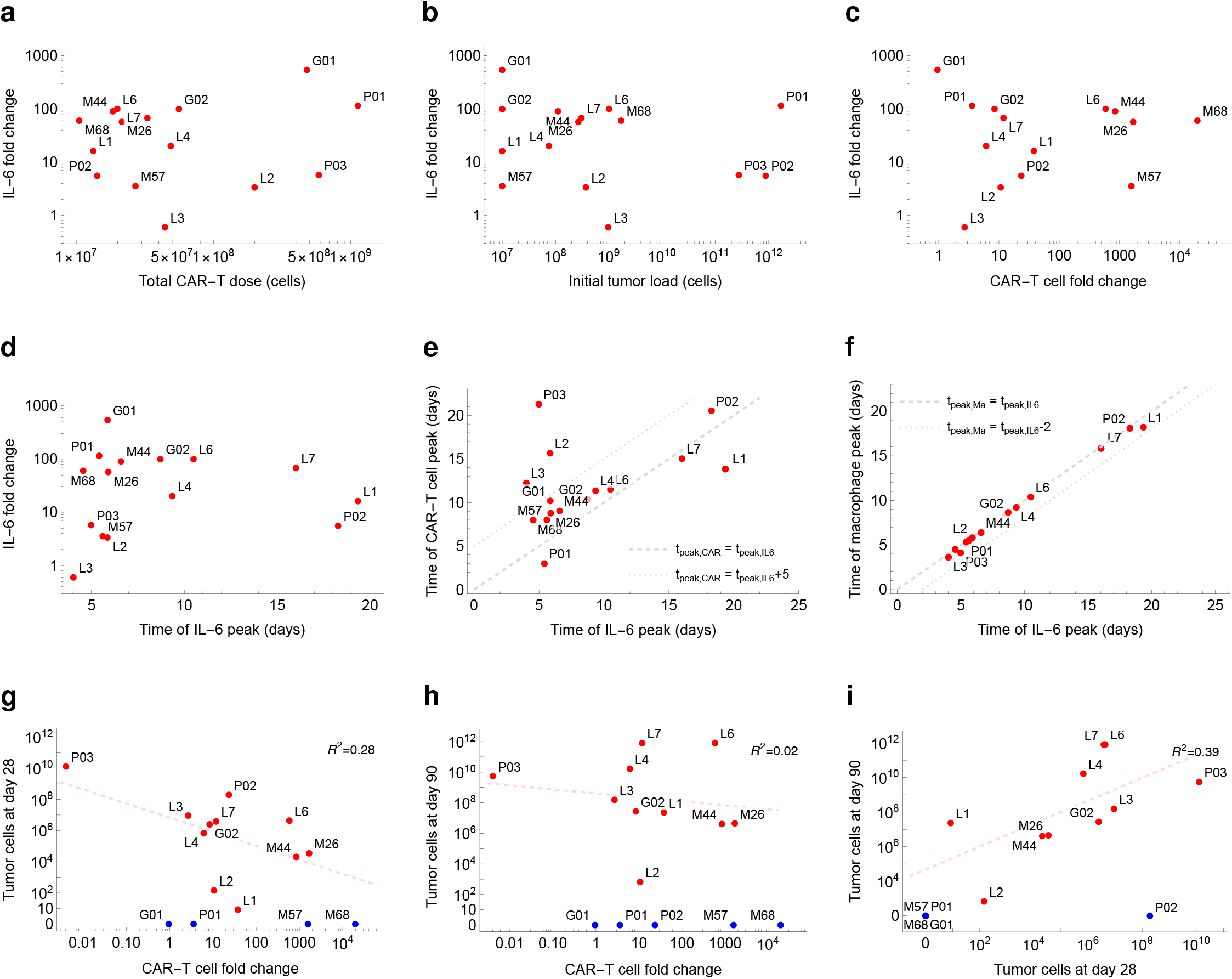
Patient-specific pharmacological and clinical parameters evaluated from model fits. **a-d** Within the patient level, the fold change in IL-6 concentration (baseline to peak) cannot be predicted from a single indicator, such as total CAR-T dose, initial tumor burden, CAR-T cell fold change (dose to peak), or time of IL-6 peak. **e** However, for the majority of patients, the model predicts that the peak of CAR-T cells occurs between 0 and 5 days after the cytokine peak (dashed lines). **f** For all patients, the cytokine peak is predicted to occur between 0 and 2 days after the peak of activated macrophages (dashed lines). **g** excluding patients predicted to present complete remission at day 28 (blue dots, patients G01, P01, M57, M68), the tumor burden at day 28 after therapy start is negatively correlated to CAR-T cell fold change. **h-i** The correlation is even weaker at day 90 since responders at day 28 become non-responders at day 90 (L4, L6, L7) while a patient with a partial response at day 28 improve its response later (P02); with exception of these cases, the model predicts an association between responses at days 28 and 90.

Our analysis revealed that, for the given cohort of 15 patients, the fold change in IL-6 concentration cannot be correlated with or predicted from the individual patient level based on clinical factors such as total CAR-T cell dose, initial tumor burden, CAR-T cell fold change, or time of IL-6 peak (Figure 5a-d). This highlights that, on the patient level, the occurrence of CRS, primarily associated with the increase of IL-6 concentrations, does not depend solely on any of the aforementioned factors but appears to emerge from an interplay of various patient-specific factors.

Although no single marker for IL-6 fold change was found, our analysis unveiled a strong temporal connection between events after CAR-T cell infusion (Figure 5e-f). The estimated CAR-T cell peaks occur within 0 to 5 days after the IL-6 peak for almost all patients, aligning with clinical observations [8]. Additionally, the cytokine peak was typically preceded by a peak of activated macrophages occurring 0 to 2 days earlier for all recorded patients. This suggests that CRS might be better predicted by monitoring the levels of activated macrophages. Although these findings need clinical validation they open a window of opportunity to intervene before a critical cytokine peak builds up, thereby lowering the risk of CRS.

We also investigated the remission level at days 28 and 90 post-therapy initiations and assessed their correlation with CAR-T cell expansion (Figure 5g-i). Neglecting the patients who achieved sustained remission, we observe a negative correlation between tumor burden and CAR-T cell fold change from dose to peak. Although we observe some correlation at day 28 (R^2^=0.28), it becomes even less clear by day 90 (R^2^=0.02). The fact that some patients shift from complete or partial remission at day 28 to different statuses at day 90 (L01, L04, L06, L07, P02) underline the complex dynamic nature of therapeutic outcomes over time. Nevertheless, the direct correlation confirms an association between responses at days 28 and 90 (R^2^=0.39).

In summary, while no single marker at the patient level reliably predicts the IL-6 peak and subsequent CRS occurrence, monitoring macrophage levels enables prediction of cytokine kinetics, thereby suggesting a potential clinical intervention.

### CAR-T cell dose and initial tumor burden influence individual treatment outcome

We showed that two crucial and clinical available parameters, namely CAR-T cell dose and initial tumor burden are insufficient to individually predict the severity of potential side effects, reflected by the poor correlation with IL-6 fold change. However, in terms of our mathematical model we can evaluate how changes of these parameters effect treatment outcome on a cohort level. Using the optimal parameter set identified for each patient and changing only the CAR-T cell dose or initial tumor burden, we study how a 10 and 100 fold change in these parameters affect typical response parameters, including CAR-T cell peak (*C*_*max*_), remission levels (*T*_*min*_) and long-term memory formation at days 28 and 90, IL-6 fold change (baseline to peak), and the time of CAR-T cell and IL-6 peaks (Figure 6, Supplementary Figure 6).

**Figure 6:**
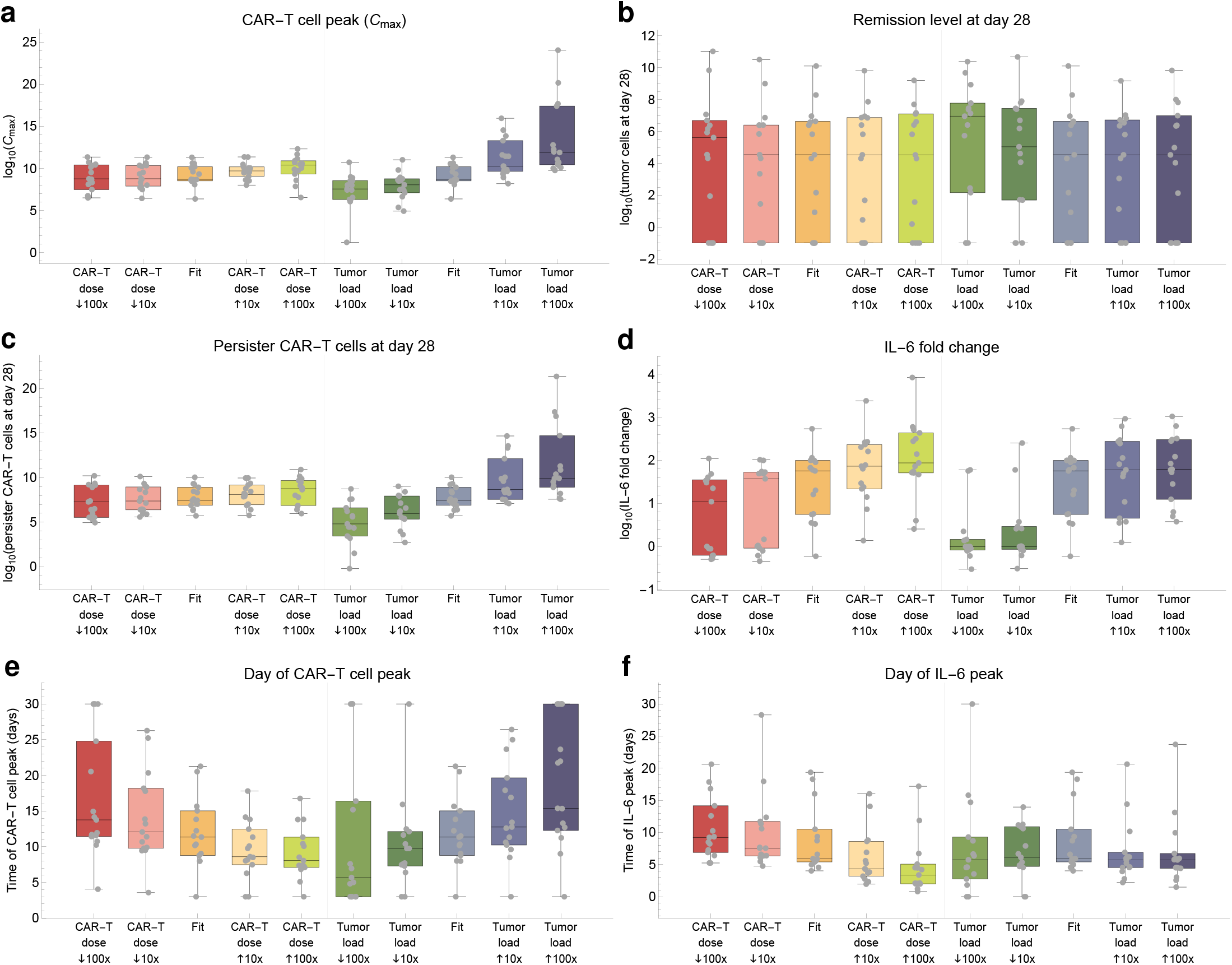
Effect of different dosing protocols on CAR-T cell dynamics, tumor response, and cytokine peak. To investigate the outcomes of a different CAR-T dose or change in the preconditioning or bridging therapy, we compared the standard scenario (Fit) with simulations starting with either a different CAR-T dose or initial tumor burden (10x and 100x smaller and higher). Assessed outcomes were: **a** number of CAR-T cells at peak (*C*_*max*_), **b** number of tumor cells at day 28, **c** number of persister CAR-T cells at day 28 **d** IL-6 fold change (baseline to peak), **e** day of CAR-T cell peak, **f** day of IL-6 peak. The number of tumor and CAR-T persister cells at day 90 were also assessed and did not present sensible differences (Supplementary Figure 6).

Upon analyzing the impact of different CAR-T cell doses, we observed that the CAR-T peak concentration, remission levels and long-term memory formation at days 28 and 90 remained almost unaffected (Figure 6a-c). However, smaller doses resulted in decreased IL-6 fold change and delayed CAR-T cell and IL-6 peaks (Figure 6d-f). This suggests that smaller CAR-T cell doses extend the expansion phase without a proportional increase in absolute cell numbers, maintaining overall remission levels. Moreover, this extended CAR-T cell expansion phase potentially leads to a more distributed activation of macrophages over time, as reflected not only in the delayed IL-6 peak but also in a decreased fold change from baseline to peak, potentially reducing the chances of severe CRS.

Initial tumor burden is an important factor that can be targeted during bridging and preconditioning therapy. We observed that reducing the initial tumor burden led to lower CAR-T cell peaks and IL-6 fold change(Figure 6a,d), with an earlier CAR-T cell peak and no substantial changes in the IL-6 peak timing (Figure 6e-f). Additionally, modifying the initial tumor burden had minimal impact on remission levels at days 28 and 90, but higher tumor burdens were associated with increased long-term memory formation (Figure 6b-c).

Although we see a correlation between IL-6 peak levels and both the CAR-T cell dose and the initial tumor burden (Figure 6d), this effect is not proportional as it would be expected from a naive approach. It rather appears that a saturation arises from the functional model setup in which antigen-binding and DAMPs serve as a trigger rather than the primary mechanism for macrophage activation (see below).

It is worth noting that important parameters regarding long-term response (namely remission level and memory formation) are not influenced by changes in the initial CAR-T dose, while a lower initial tumor burden does not affect remission level but leads to less memory formation. Moreover, side effects due to cytokine release could potentially decrease with smaller administered CAR-T cell doses and intensive pre-conditioning to reduce the initial tumor burden.

In conclusion, our findings suggest that adjusting the administered CAR-T cell dose may exert a greater influence on kinetic parameters such as peak times and IL-6 peak concentration. The initial tumor burden and the corresponding antigen load appears to serve as a catalyst for CAR-T cell expansion, potentially explaining higher persistence of CAR-T cells but also increased severity of cytokine-related side effects.

### Disentangling mechanisms of macrophage activation and CRS timeline

While it appears that cytokine-related side effects can be reduced by adjusting CAR-T cell dose or initial tumor burden, targeting macrophage-mediated cytokine release directly is also a clinical option. A quantitative understanding of the temporal dynamics of macrophage activation and cytokine release is crucial for this purpose.

Our model assumes three distinct molecular and cellular mechanisms for macrophage activation, namely DAMPs release, antigen-binding, and CD40 contact, described by parameters *β*_*K*_, *β*_*B*_ and *β*_*C*_ respectively (equation (11)). To unveil the dynamics underlying these mechanisms, we simulated the time course of patient M44, selectively activating one mechanism at a time (Figure 7a). Activation via DAMPs release alone (*β*_*K*_ *>* 0, *β*_*B*_, *β*_*C*_ = 0) led to IL-6 peak around day 3, during the tumor shrinkage phase. Activation solely through antigen recognition (*β*_*B*_ *>* 0, *β*_*K*_, *β*_*C*_ = 0) resulted in IL-6 concentration rising concurrently with CAR-T cell expansion, peaking around day 8. Considering only CD40-mediated activation (*β*_*C*_ *>* 0, *β*_*K*_, *β*_*B*_ = 0), IL-6 kinetics remained at baseline levels due to the requirement of previous infiltration by naive macrophages. Yet, integrating CD40-mediated activation with small values for DAMPs release and antigen-binding parameters (*β*_*K*_ *>>* 0, *β*_*B*_, *β*_*C*_ ≈ 0), the IL-6 peak occurred around day 13, four days post-CAR-T cell peak (purple curve). These findings suggest that each mechanism occurs at distinct time points, influenced by various interactions unfolding after infusion: DAMPs drive macrophage activation during the tumor shrinkage phase, followed by antigen-binding-mediated activation during CAR-T cell expansion, and contact-dependent CD40-mediated activation.

**Figure 7:**
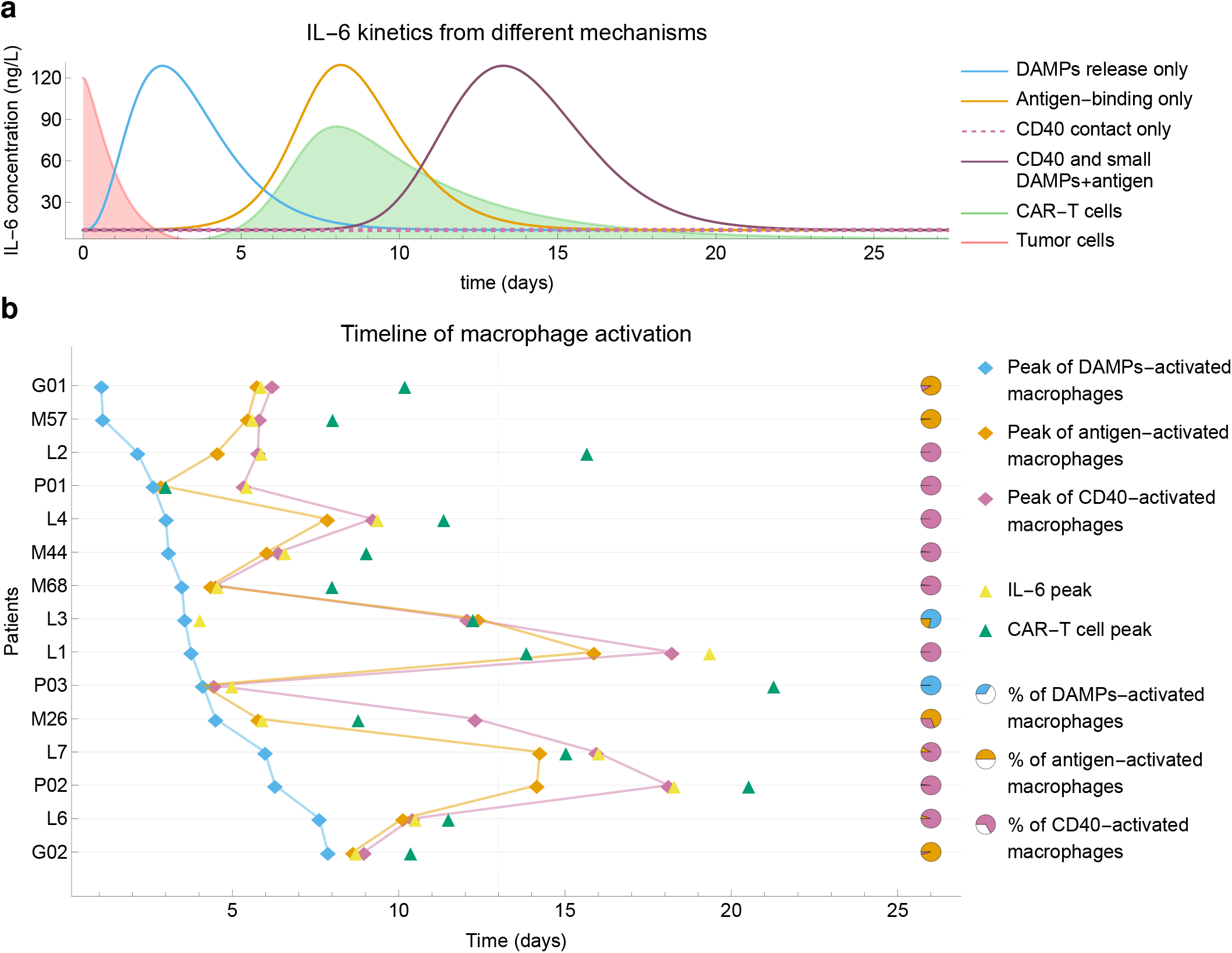
Dynamics of macrophage activation and cytokine release. **a** Time course of patient M44, selectively activating DAMPs release, antigen-binding, and CD40 contact mechanisms one at a time. DAMPs-only activation leads to an IL-6 peak around day 3 during tumor shrinkage. Antigen-biding activation results in IL-6 increasing with CAR-T expansion, peaking around day 8. CD40-mediated activation alone maintains IL-6 at baseline, but when combined with small values for DAMPs and antigen-binding parameters, it produces an IL-6 peak around day 13, four days after the CAR-T cell peak. **b** Splitting the active macrophage population according to their source, i.e., activation mechanisms (see Methods), and reconstructing the timeline for each patient, we found that the peak of DAMP-activated macrophages takes place early, during the tumor shrinkage phase. The second sub-population to present a peak is the antigen-binding-activated macrophages, which expand in parallel with CAR-T expansion. The last sub-population to peak are CD40-activated macrophages, which require the presence of previously activated macrophages. For 8 out of the 15 patients, the major source of activated macrophages was CD40 contact (pie charts), which was also the source of 62% of all activated macrophages when all patients are combined.

To understand the individual contributions of each mechanism, we divided the activated macrophages into three sub-populations based on their source (DAMPs-activated, antigen-activated, CD40-activated) and, for each patient, determined the peak time and cumulative number of activated macrophages for each sub-population (Supplementary Text S5), and the CAR-T cells and IL-6 peak times (Figure 7b). Interestingly, the previously described timeline holds true for the majority of patients: DAMP-activated macrophages reach their peak levels first, followed by the peak of antigen-activated macrophages, followed by the peak of CD40-activated macrophages.

While all these mechanisms can in principle equally contribute to macrophage activation, a computational estimate of the cumulative number of activated macrophages in each sub-population revealed that the majority of activated macrophages (62% considering all patients) are activated by the CD40 contact-dependent mechanism (pie charts in Figure 7b). This intriguing discovery suggests that macrophage activation mediated by contact-independent mechanisms, such as DAMPs release and the release of GM-CSF and IFN-*γ* by antigen-binding-activated CAR-T cells, is limited and may not be the primary source of inflammation and CRS. On the contrary, our findings propose that once CD40-mediated activation is triggered by these previous mechanisms, it enters a feedback loop in which activated macrophages in contact with CAR-T cells further activate more macrophages, leading to massive cytokine release. These results confirm that CD40–CD40L interactions are not essential for eliciting cytokine release but are likely the primary drivers exacerbating macrophage activation, IL-6 production, and CRS severity. To address the robustness of these results, an extensive analysis comparing the full model with its reduced versions considering one or two mechanisms only confirmed our findings (Supplementary Text S6 and Supplementary Figures 7 and 8).

In summary, our findings indicate that macrophage activation via DAMPs release, antigen-binding, and CD40 contact unfolds sequentially during therapy, with CD40-mediated inflammation playing a predominant role.

### Targeted interventions to control macrophage activation

The deconvolution of the different mechanisms driving macrophage activation hints towards a clinical intervention to mitigate CRS. Currently, the primary drug used to address CRS is Tocilizumab, a monoclonal antibody that acts as an IL-6 antagonist [12]. However, targeting macrophage activation with GS-CSF and CD40 antibodies has also been explored in cell-based studies [42].

To mimic the use of such antibodies that specifically inhibit macrophage activation, we applied our patient-specific model but reduced the parameters related to macrophage activation, *β*_*K*_, *β*_*B*_ and *β*_*C*_ one at time (Figure 8a). We observe that the IL-6 peak decreased with increased blocking, with varying effects depending on the patient. Overall, for a 50% reduction in the respective activation parameter, the mean reduction in the IL-6 peak was 7%, 12%, and 34% for DAMP, antigen-binding, and CD40, respectively. Repeating the same simulations for reduced, alternative models showed a similar or greater reduction in the IL6 peak when CD40 activation is reduced by 50% (Supplementary Figure 8). In addition, sensitivity analysis showed that these values did not change substantially when each of the eight fitted parameters was varied by 50% (Supplementary Figure 9). This suggests that the CD40-CD40L interaction is not only a major driver of CRS but also a promising target for clinical interventions.

**Figure 8:**
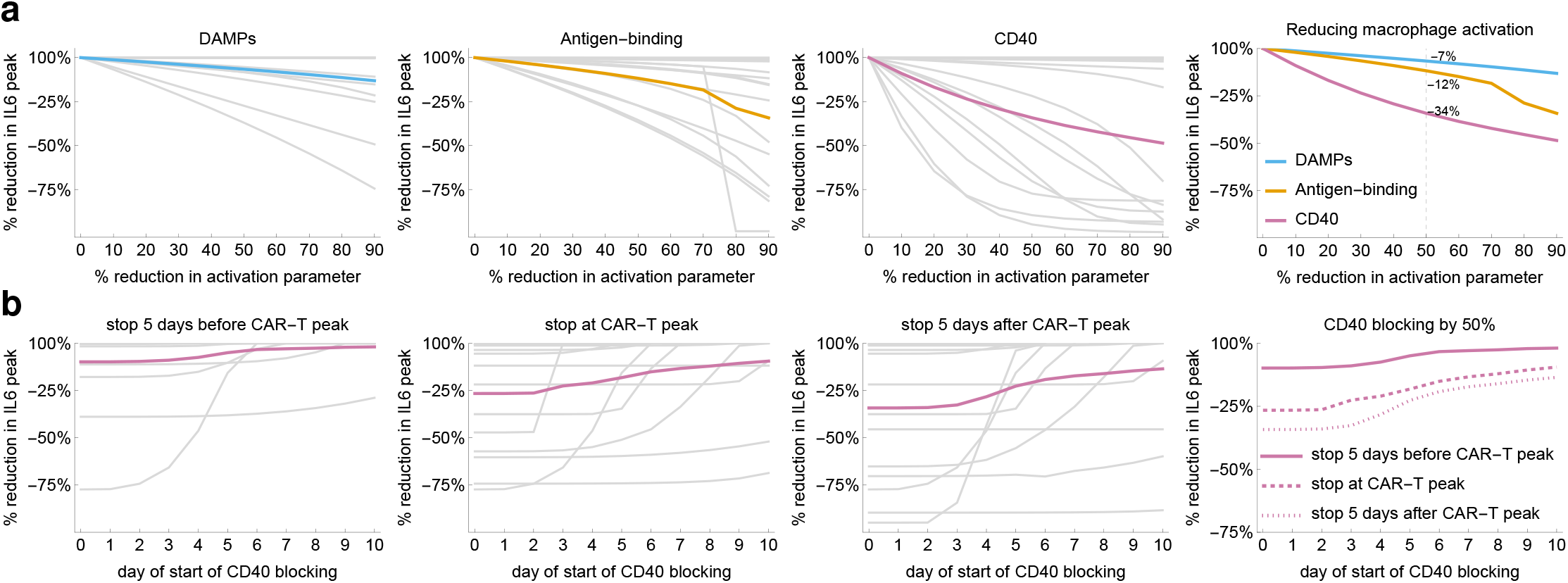
Blocking of macrophage activation and cytokine release. **a** Simulation results of interventions reducing mechanisms of macrophage activation one at a time, showing the reduction in IL-6 peak for each patient (gray curves) and mean effect (colored curves). For a simulated 50% reduction (dashed line, fourth panel) the overall reduction in IL-6 peak is indicated for each mechanism. **b** Simulation results of interventions blocking CD40 starting 0-10 days after CAR-T cell infusion, stopping -5, 0 and 5 days after CAR-T cell peak. The plots show the resulting reduction in IL-6 peak for each patient (gray curves) and the mean effect (colored curves).

To demonstrate the clinical potential of CD40 blocking, we conducted a series of model simulations reducing, for each patient, the CD40 parameter to 50% of the original value starting at different time points after infusion and keeping the block until either before, at or after the CAR-T cell peak (Figure 8b). We found that initiating the blocking 1 or 3 days after CAR-T cell infusion is most effective, with the block being maintained until or after the CAR-T cell peak. On average, a 30-40% reduction in IL-6 peak can be achieved with some patients showing even stronger responses.

Our model of sequential macrophage activation suggests that targeting the CD40-CD40L axis is an effective and clinically feasible strategy to control macrophage activation and reduce IL-6 peak height.

## Discussion

We developed a novel multi-layer mathematical model to investigate tumor response and CRS following CAR-T cell immunotherapy. The model accurately describes the timeline of therapy response and relates its dynamic shape to interpretable model parameters. Our findings highlight the role of early macrophage activation in cytokine release, thereby outlining potential strategies for clinical monitoring and prevention of CRS. By disentangling different modes of macrophage activation we identified the CD40-CD40L axis as a clinically feasible target to control the activation process and modulate IL-6 peak height.

The diverse configuration and clinical outcomes of patient responses to CAR-T cell therapy is attributed to product- and patient-specific factors [39]. Our model unveiled the mechanisms underlying multiphasic therapy response, showing how each of its shape features is linked to a specific model parameter. While, on one hand, this close correlation is the basis for a model-informed step-wise parameter estimation strategy, it, on the other hand, explains how key clinical indicators (CAR-T cell peak, persistence, and minimal tumor burden) are modulated by product-related factors (CAR-T cell expansion, cytotoxicity and receptor density) and patient-specific variables (tumor growth rate and antigen density).

In our analysis of patient cohorts, neither the initial CAR-T cell dose nor the baseline tumor burden proved to be reliable predictors for treatment outcomes. These factors, along with maximum CAR-T cell concentration, CD4/CD8 ratio, pre-infusion phenotypes, area under the concentration-time curve (AUC), have been extensively studied but lack consensus as prognostic factors [43, 44, 38, 45, 27]. On the other hand, we found that the minimal tumor burden is inversely related to the CAR-T cell peak concentration, consistent with [46], which reported that higher peaks of CAR-T cells within 15 days are associated with increased likelihood of achieving complete remission.

While the precise determinants of CRS severity remain debated, some studies found tumor burden, lymphodepletion, CAR-T cell dose, and CAR-T product are independent predictors of CRS [47, 48] and others suggested that initial tumor burden and CAR-T cell dose correlate with the severity of CRS [49, 50, 51]. In our patient-level analysis, neither CAR-T cell dose nor initial tumor burden reliably predicted IL-6 increase (as a surrogate for CRS severity) a priori. However, our cohort simulations shows that smaller CAR-T cell doses reduced IL-6 fold change, potentially lowering severe CRS risk without sensitive affecting CAR-T cell peak, remission levels, or memory formation. These findings aligns with the previously reported association between the expansion of CAR-T cells (peak/dose) and severity of CRS [43]. Also, a lower initial tumor burden reduced IL-6 levels but lowered both CAR-T cell peak and persistence, with minimal impact on remission levels. In [52], markers related to tumor burden and inflammation, both of which may be influenced by the underlying tumor biology, were highly associated with clinical outcomes.

Several mathematical models have been developed to understand T cell and cytokine interactions in different contexts [53, 54, 55, 56]. For CAR-T cell therapy, Mostolizadeh et.al [23] developed a model that includes CAR-T cells, healthy and cancer cells, and cytokines, and applied optimal control theory for controlling cytokine release syndrome with tocilizumab. Hardiansyah and Ng [22] implemented a quantitative systems pharmacology model that links CAR-T cell kinetics to cytokine release, showing a correlation between disease burden and CAR-T cell expansion. Zhang et al. [25] built a computational model of CRS during CAR-T cell therapy, illustrating how cellular and molecular interactions among CAR-T cells, B-ALL cells, bystander monocytes, and various inflammatory cytokines affect CRS severity. In [24], the modulation of drug administration frequency and duration to maintain efficacy while preventing cytokine storms was investigated. While some of these models have been validated with clinical data, none explicitly include how different macrophage-associated mechanisms influence the extent and temporal dynamics of CRS.

Mouse models showed that CRS involves a multicellular network comprising CAR-T and host cells, with macrophages playing a crucial role [14, 16]. A recent review [8] proposes that macrophage-mediated CRS emerges from antigen-binding-mediated inflammation, DAMPs release and CD40 contact. Built upon these concepts, our model accurately replicates these dynamics, showing an early peak of activated macrophages followed by cytokine peak within 1-2 days, then a CAR-T cell peak within 1-5 days. Disentangling these mechanisms, we showed that DAMPs underlie early cytokine release, while antigen-binding drives cytokine increase parallel to CAR-T cell expansion, and CD40 contact leads to late cytokine rise. A deeper analysis indicates that CD40 is the latest but the main driver of macrophage activation, suggesting that antigen-binding and DAMPs act more as triggers than primary mechanisms.

The preponderant role of CD40-contact in macrophage activation poses a candidate for interventions against CRS. Simulating a reduction of the different activation mechanism showed that targeting CD40 leads to the most substantial reduction in IL-6 peak height, causing an average reduction of 34% in the case of constant blocking and 33% reduction when blocking from day 3 after infusion to 5 days after CAR-T cell peak. This aligns with a recent cell-based in vitro model for CRS that showed that IL-6 supernatant levels decreased 20-30% when blocking cytokine release by bystander macrophages with neutralizing antibodies for GM-CSF and CD40, and decreased 30–40% due to the genetic disruption of the CD40L and/or CSF2 knock-out CAR-T cells [42].

Our model and corresponding results are inherently limited by the underlying and simplified concept of CAR-T cell development, which assumes a progression from naive to expander phenotype and then to a persister phenotype. This concept is consistent with the classical view of T cell biology [57] and long-standing models of T cell responses [58, 59, 26]. It also builds on a recently proposed framework for mechanistic models of CAR-T cell dynamics [36] and previous work in the field [38, 60, 5, 61, 7]. However, alternative pathways for T cell differentiation from naive to memory and then to effector cells have been proposed [37]. Although there is recent evidence that some types of memory cells have the ability to expand [62], it has also been shown that in models in which only long-term persistent cells are able to expand, the expected T cell kinetics cannot be described with realistic parameter values [57]. Therefore, we adhere to a widely accepted view of T cell biology, by considering the naive-expander-persister development. Moreover, following [36], our population of CAR-T expanders can be viewed as a composition of effector cells and eventually expanding memory cells (stem, central and effector memory), while the CAR-T persister cells comprise the long-lived memory cells. Finally, the inclusion of a bidirectional, antigen-dependent phenotypic switch between these two populations further implies a plasticity-like development rather than a unidirectional differentiation pathway. Other pathways of CAR-T cell development or mixed phenotypes for the initial conditions were not considered here for parsimonious reasons. As a consequence, our model is close to minimal with respect to the number of parameters. This is highlighted by the correspondence between each model parameter and a shape feature of the multiphasic CAR-T and tumor cell responses. In conclusion, T-cell development is a complex and multifactorial process, and it remains to be elucidated to what extent specific differentiation pathways are taken and how much quantitative results on CAR-T cell treatment depend on these aspects.

The nature of our research was investigative rather than predictive, aiming to understand the mechanisms of therapy response and macrophage-mediated CRS rather than making predictions at a populational level. While our model provides insights into these processes, it also faces limitations. The one-to-one correspondence between parameters and time course shape features allows comprehensive understanding of system dynamics and partially addresses the challenge of parameter identifiability. However, this process requires tumor data at several time points, which is not available in most cases [31, 1, 32, 33, 34]. Moreover, although preclinical models indicated that IL-6 plays a role in macrophage activation, the exact degree of this contribution remains uncertain [14]. Due to the need for additional data to accurately calibrate this relationship, we opted not to incorporate a feedback mechanism of IL-6 on macrophage activation, regarding it as a second-order effect. In addition, the use of macrophage time courses may improve our fitting strategy, which estimates IL-6 and macrophage parameters based on IL-6 data only. Finally, while we use IL-6 fold increase to indicate CRS severity, there is no standard-ization in clinical grading scales [63], which are often based on symptoms or therapeutic interventions rather than measurable thresholds [64, 65, 66].

Our study demonstrates the efficacy of a compact multi-module mathematical model in accurately describing a retrospective dataset from patients undergoing CAR-T cell therapy. Our findings shed light on the timeline and influence of macrophage activation on cytokine release and pinpoint the CD40-mediated mechanism as the primary driver of cytokine release, suggesting it as a potential target for CRS treatment strategies.

## Supplementary material

Supplementary material contains Supplementary Texts S1–S6, Supplementary Figures 1–11 and Supplementary Tables 1–3.

## Data availability

The patient data that support the findings of this study were collected from existing literature [1, 31, 32, 33, 34].

## Author contributions

Conceptualization: DS, LRCB, IG, ACF. Formal Analysis, investigation, methodology: DS, ACF. Writing: DS, LRCB, IG, ACF. Supervision: ACF.

