## Supplementary Material for "Mathematical modeling unveils the timeline of CAR-T cell therapy and macrophage-mediated cytokine release syndrome"

### Supplementary Texts

#### S1. Model setup

Here we describe details of model setup, initial conditions, and unit conversion. In instances where the initial tumor burden was either unspecified (NA) or had 0% blasts, we established a baseline of $T_P(0) = 10^7$ cells. Data points where %blasts were approximately 0 after therapy were not included in calibration due to the high uncertainty associated with digitization. For patients in the studies by Ma et al. [1] and Li et al. [2], where tumor burdens were expressed as percentages, we made an approximation based on the assumption that this percentages are related to white blood cells (WBC) and an adult has 5L of blood. Using an WBC range of $(2.0 - 6.5) \times 10^{10}$ provided in [3], we have an average of $4.25 \times 10^{10}$ of WBC cells. The absolute number of tumor cells was then estimated using the formula: tumor cells = WBC cells × % blasts.

Despite an extensive literature search for datasets with patient time courses sufficient for CAR-T multiphase and IL-6 kinetics, there was insufficient data to describe the phenotypic characterization of the CAR-T product upon infusion. Consequently, we assumed the administered dose of CAR-T cells as the initial condition of the injected CAR-T cell population ($C_I(0)$). The initial populations of other phenotypes were set to zero ($C_E(0) = C_P(0) = 0$) as these phenotypic changes are driven by antigen-binding and occur throughout the dynamics' evolution. For antigen-negative tumor cells, except for patient G02, the initial tumor burden ($T_N(0)$) was fitted below the detection limit of 2.5$\times 10^5$ cells.

The scaling factor for CAR-T cell count data was estimated using data provided by Lee et al. [4] and Kalos et al. [5], where both counts per microgram and total circulating cells were reported. The conversion factor was determined to be 1 CAR-T cell copy/$\mu$g DNA = $10^4$ CAR-T cells. For conversions between the absolute number of CAR-T cells and cells/$\mu$L, we used the same blood volume of 5L in humans, which gives us 1 CAR-T cell/$\mu$L = $5 \times 10^6$ CAR-T cells. Furthermore, considering that about 1% of cells are present in the peripheral blood (PB) at all times compared to the bone marrow (BM), we have 1 CAR-T cell/$\mu$L in BM = $5 \times 10^8$ CAR-T cells in PB.

To prevent artificial regrowth, we applied zero-limits to CAR-T and tumor cell populations: if any cell population reached less than one cell, proliferation was set to zero.

#### S2. Parameter estimation for CAR-T and tumor cells

A brief assessment of the model multi-layer structure shows that a two-step fitting procedure is equivalent to a one-step approach that fits all model parameters at once. Indeed, since equations (1-7)

for CAR-T and tumor cells are independent of macrophages or IL-6, writing the optimal fit as $(p_1, p_2)$, where $p_1$ contains the CAR-T and tumor cell parameters while $p_2$ contains the macrophage and IL-6 parameters, we see that varying $p_2$ does not change the equations (and thus the residuals) of the first layer. Thus, obtaining a global fit $(p_1, p_2)$ is equivalent to first optimizing $p_1$ for the CAR-T and tumor cell dynamics (equations (1-7)) and then optimizing $p_2$ to fit IL6 dynamics (equations (8-11)).

Therefore, for all 25 patients with CAR-T and tumor cell time courses, we first estimated the 12 parameters for the CAR-T and tumor multiphasic dynamics (Figure 1a,b).

First, to reduce the number of free, patient-specific parameters, we analyzed the parameters that least influenced the multiphasic dynamics of CAR-T and tumor cells, and defined constant, universal values for parameters $K$ (carrying capacity of tumor cells), $\theta$ (recruitment rate of persister CAR-T cells). We then defined reasonable ranges for the other 10 parameters, using literature data when available, see Supplementary Table 1.

Second, we performed a manual and iterative process to qualitatively fit the remaining 10 patient-specific parameters. This process was based on the analysis that identified a mapping between the 10 shape features and the 10 patient-specific parameters (see Figure 4) and is described as follows. We first analyzed the overall CAR-T cell dynamics on a logarithmic scale, segmenting the data into distinct phases: distribution, expansion, contraction, and persistence. For each phase, we determined an exponential curve, establishing initial estimates and bounds for the dominant parameters ($\mu_I, \mu_E, \mu_P, \kappa$), see Supplementary Figure 10 and Supplementary Table 1. Combining these initial approximations with our knowledge of the influence of the other parameters ($\eta, \epsilon, A, B, \gamma, \rho$) in each shape feature of the model solutions (Figure 4), we manually obtained the initial estimates for the 10 patient-specific parameter values.

Finally, these initial estimates were further improved by an appropriate optimization algorithm that guarantees local minima. To perform this step, we encoded the model structure in QSP Designer [6] and employed the Nelder-Mead method for parameter estimation, minimizing the weighted least squared error (WLS) between model simulations and data. Weights were defined as the inverse of the square of the maximum observed values. After initial runs, we refined the parameter ranges, and this iterative process continued until an adequate fit to the overall CAR-T cell dynamics was achieved.

The same approach was applied to the extended model for patients with antigen-negative tumor relapse, equations (6-7), with an extra free parameter $g_0$ (fraction of citotoxicity reduction for antigen-negative tumor cells), which we estimated to be a small fraction [7].

With this process, we obtained the fits for CAR-T and tumor cells shown in Figure 2 and Supplementary Figures 1, 2, 3. These values are provided in Supplementary Table 3.

### S3. Parameter estimation for IL-6 dynamics

After fitting the model parameters for the first layer encompassing CAR-T and tumor cells, we fitted the macrophage and IL-6 parameters using IL-6 time-courses of 15 patients.

Equations (8-11) have 12 parameters, namely $\sigma_M, \delta_M, \sigma_I, \delta_I, \alpha, \beta_B, \beta_K, \beta_C, C, M_i(0), M_a(0)$ and $IL_6(0)$. After initial analysis of the model dynamics using parameter values in reasonable ranges, we fixed the following 4 parameters as follows. The initial number of activated macrophages was set to zero, $M_a(0) = 0$. The initial condition for IL6 was obtained from the first time point for each patient, $IL_6(0) = IL_{61}$. In absence of IL-6 release by activated macrophages, IL-6 levels reach the steady state $\sigma_I/\delta_I$, which is therefore a lower bound for the minimum IL-6 level during the response phase, thus we set $\sigma_I = \delta_I \min_i IL_{6i}$ for each patient. The model dynamics did not present high sensitivity to saturation parameter $C$, so it was also fixed; since the range for the initial number of inactive macrophages $M_i(0)$ was estimated between $[10^9, 10^{11}]$ cells [8, 9, 10], we set the value $C = 10^{10}$ to describe a saturation within the range of inactive macrophages. For the 8 remaining parameters we defined biologically reasonable ranges based either on literature estimates or steady state conditions, with values provided in

Supplementary Table 1.

Using the defined parameter ranges, we applied an automated routine coded in *Mathematica* and based on the strategy used in [11], combining global and local optimization methods to estimate patient-specific parameter values. First, for each patient, it performs a global search using a Monte Carlo approach, consisting of simulating the model for $10^4$ randomly chosen parameter tuples within these ranges. For each simulation, the logarithm of the weighted sum of squares is computed as

$$LR = \log\left(\sum_{i=1}^{n_d} w_i \left(IL_6(t_i) - IL_{6i}\right)^2\right), \tag{1}$$

where $(t_i, IL_{6i})$ are the $n_d$ individual time points. To capture the IL-6 peaks, we used weights given by $w_i = w_0 + (1 - w_0)IL_{6i}/\max_j(IL_{6j})$. A value $w_0 = 0.15$ was chosen after initial tests, giving a weight approximately seven times higher to the IL-6 peak in comparison with the minimum value of $IL_{6i}$.

The routine then selects the 100 parameter tuples that give the minimum $LR$ and refines these estimates by applying to each one a local minimization approach. The best fit is then selected as the one that gave the minimum $LR$. Using this approach, we obtained the fits shown in Figure 2 and Supplementary Figure 4, which were used in the Results Section.

### S4. Approximating solutions and predicted slopes for multiphasic dynamics

Here, we present the rationale that allows to simplify the ODEs for total CAR-T and tumor cells. This simplification leads to the calculation of approximate solutions for each phase, along with their characteristic slopes. These arguments are supported by a quantitative approach in which, for each patient, we calculated the characteristic slopes from the model solution at each phase and compared them with the predicted parameter governing each slope (Figure 3).

The total CAR-T cell population is given by $C = C_I + C_E + C_P$ and is described by

$$\frac{dC}{dt} = -\underbrace{\mu_I C_I}_{\text{Injected death}} + \underbrace{\kappa F(T_P) C_E}_{\text{Expander proliferation}} - \underbrace{\mu_E C_E}_{\text{Expander depletion}} - \underbrace{\mu_P C_P}_{\text{Persister death}}.$$

During the distribution phase, the CAR-T cell population consists mainly of injected cells, from which we approximate $C \approx C_I$ and $C_E, C_P \approx 0$. Thus, $dC/dt \approx -\mu_I C$, leading to an approximating solution $C = C_0 e^{-\mu_I t}$ with slope $-\mu_I$. During the expansion phase, the tumor burden is high and the CAR-T cell population consists mainly of CAR-T expanders, leading to the approximation $C \approx C_E$, $C_I, C_P \approx 0$, $F(T_P) \approx 1$. The approximating ODE is $dC/dt \approx (\kappa - \mu_E)C$, leading to an approximating solution $C = C_{\text{min}} e^{(\kappa - \mu_E)t}$ with slope $\kappa - \mu_E$. Similarly, in the contraction phase the CAR-T cell population presents the same characteristics, but with a reduced tumor burden, leading to $F(T_P) \approx 0$ and $dC/dt \approx -\mu_E C$. The approximating solution reads as $C = C_{\text{max}} e^{-\mu_E t}$ with slope $-\mu_E$. Finally, the persistence phase is characterized by the presence of CAR-T persisters, leading to $C \approx C_P$, $C_E, C_I \approx 0$ and an approximating solution $C = C_{\text{con}} e^{-\mu_P t}$ with slope $-\mu_P$.

Similarly, the total tumor population, given by $T = T_P + T_N$, is described by

$$\frac{dT}{dt} = \underbrace{\rho T\left(1 - \frac{T}{K}\right)}_{\text{Tumor growth}} - \underbrace{\gamma \frac{C_E}{B + C_E}(T_P + g_0 T_N)}_{\text{CAR-T killing}}.$$

During the transient phase, the number of effector CAR-T cells is small and, due to the preconditioning therapy, the tumor burden is far below the carrying capacity. Also, $C_E \approx 0$, and the dynamics is approximated by $dT/dt \approx \rho T$, leading to an approximating solution $T = T_0 e^{\rho t}$ with slope $\rho$. During the shrinkage phase, the number of CAR-T cells is very high, leading to $C_E/(B + C_E) \approx 1$ while the majority (if not all) of tumor cells is antigen-positive, $T \approx T_P + g_0 T_N$. Then, $dT/dt \approx \rho T - \gamma T$ and the approximating solution is $T = T_{\text{shr}} e^{(\rho-\gamma)t}$ with slope $\rho - \gamma$. Finally, if the tumor cells were not extinct, the tumor response phase starts during the persistence phase of CAR-T cells with small number of effector CAR-T cells. Therefore, $dT/dt \approx \rho T$ and the approximating solution is $T = T_{\text{min}} e^{\rho t}$ with slope $\rho$.

### S5. Assessing different sources of macrophage activation

To asses the different sources of macrophage activation, we split the macrophage population in three sub-populations according to the activation mechanism, $M_a = M_{a,K} + M_{a,B} + M_{a,C}$, where $M_{a,K}$ are the DAMP-activated macrophages, $M_{a,B}$ are the antigen-binding-activated macrophages and $M_{a,C}$ are the CD40-activated macrophages. The differential equations for each population are obtained by splitting the activation rate (11), leading to

$$\frac{dM_{a,j}}{dt} = h_j M_i - \delta_M M_{a,j}, \quad j = K, B, C, \tag{2}$$

where the activation of each sub-population is given by

$$h_B = \beta_B \frac{T_P}{A + T_P} C_E, \quad h_K = \beta_K \frac{C_E}{B + C_E}(T_P + g_0 T_N), \quad h_C = \beta_C \frac{M_a}{C + M_a} C_E. \tag{3}$$

With this, we calculated for the simulated solutions the peak time for each macrophage population. Further, the total number of macrophages activated by each mechanism is obtained by evaluating the integrals $\int_0^{t_F} h_j M_i dt$, $j = K, B, C$ until a final time $t_F$.

### S6. Model comparison

To test the robustness of our conclusions on the timeline of macrophage activation, and to determine the relative importance of each mechanism in driving IL-6 dynamics, we compared the full model with alternative, reduced model structures. Denoting the model mechanisms as D (DAMPs release), A (antigen-binding) and C (CD40 contact), we compared the full model (denoted DAC model) with the reduced models DC, AC, DA, D and A, each corresponding to the removal of one or two mechanisms at a time and corresponding to setting $\beta_K$, $\beta_B$ or $\beta_C$ to zero. The model with CD40 contact only (model C) was not tested as it does not result in macrophage activation at all since the CD40 axis requires the presence of previously activated macrophages (see equation (9) with $\beta_K = \beta_B = 0$). For each alternative model, we repeated the same automated parameter estimation procedure described above and determined the best fit for each patient. For each model and each patient $j$, we calculated the logarithm of residual sum of squares $LR_j$ (equation (1)) and the AIC criteria, given by

$$\text{AIC}_j = 2k + n_{d,j}(1 + LR_j - \ln n_{d,j} + \ln 2\pi), \tag{4}$$

where $k \in \{6, 7, 8\}$ is the number of free parameters and $n_{d,j}$ is the number of data points of patient $j$. The full DAC model had the minimum sum of $LR_j$ and $AIC_j$ and was the best model for 11 out of 15 patients, followed by models DC in second place and AC in third, both with $LR_j$ and $AIC_j$ in the same range (Supplementary Figure 7). Interestingly, both AC and DC models assume CD40 contact as an activation mechanism, differing only on the triggering mechanisms (antigen-binding in the AC model and DAMPs release in the DC model). The analysis performed for the DAC model in Figure 7 was repeated for the reduced AC and DC models and resulted in the same timeline for macrophage activation as well as an equal or higher percentage of CD40 activated macrophages (Supplementary Figure 8a,c). Simulating a 50% reduction in each activation mechanism also led to similar reductions in IL-6 peak (compare Figure 8a and Supplementary Figure 8b,d). These results show that at least two mechanisms are required to accurately describe the dynamics of IL-6, with CD40 contact implied to be the main driver of CRS.

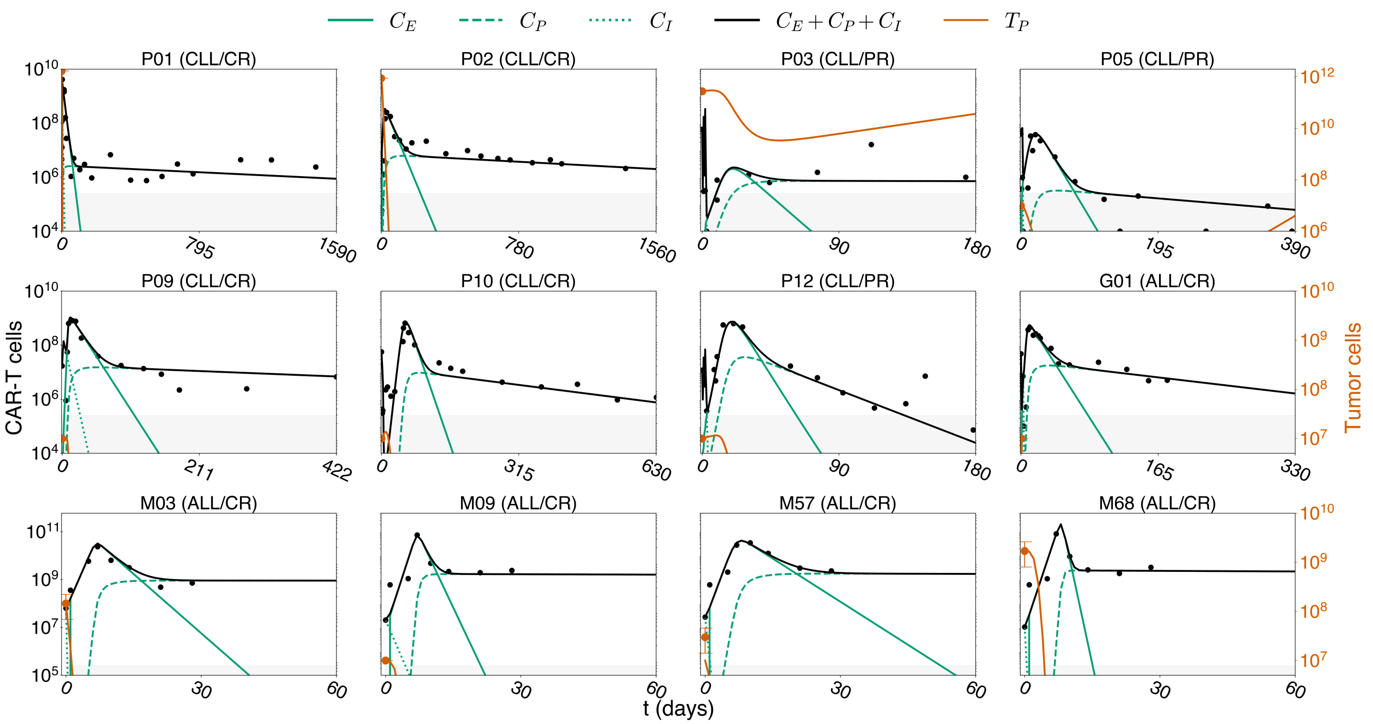


**Supplementary Figure 1. Model simulations for responders.** Model fits for selected patients that showed either complete response (CR) or partial response (PR). CR is achieved when the tumor is clinically undetectable, while PR is defined by a final tumor burden at least less than 50% of the initial burden. The CAR-T cell detection threshold of $2.5 \times 10^5$ cells is represented by the gray shaded area.

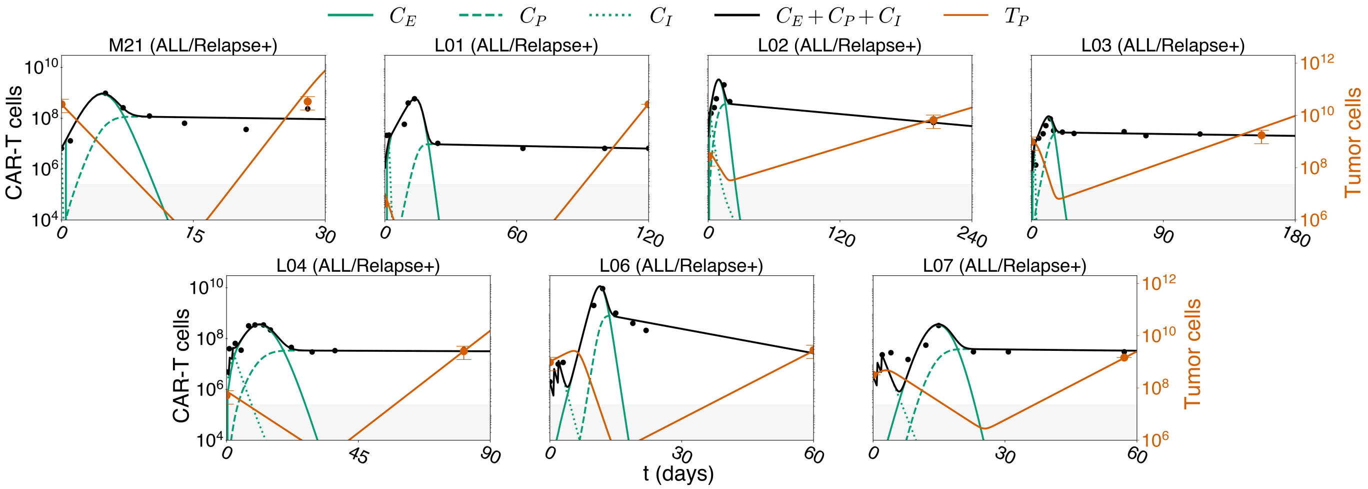


**Supplementary Figure 2. Model simulations for patients with antigen-positive relapse.** Model fits for selected patients that showed relapse of antigen-positive tumor cells, excluding data points where %blasts approached zero. Tumor cell error bars represent the range of WBCs used in scaling (see Methods). The CAR-T cell detection threshold of $2.5 \times 10^5$ cells is represented by the gray shaded area.

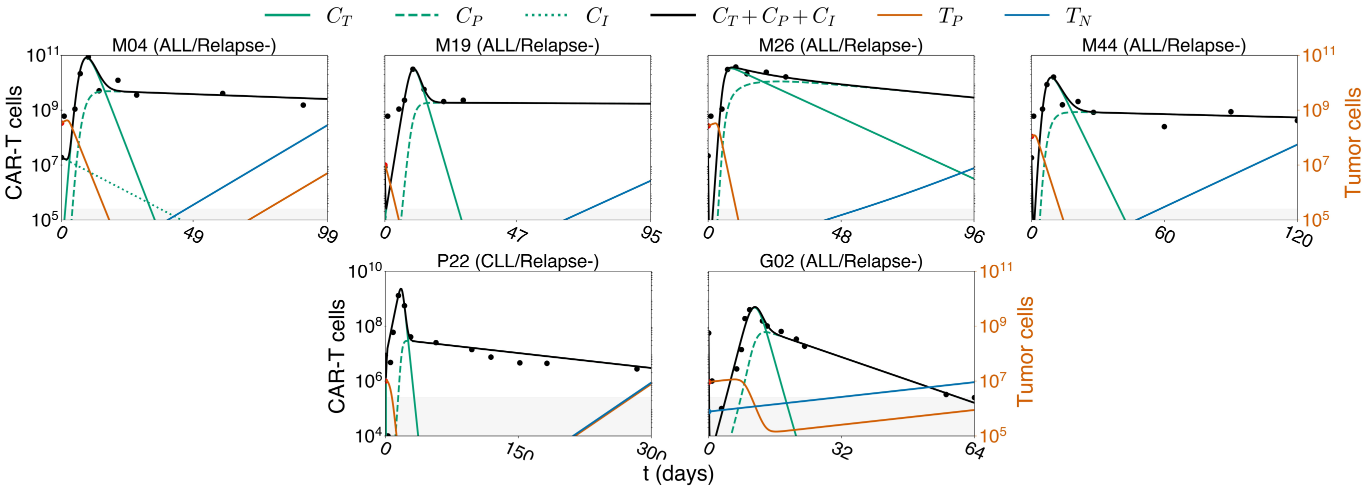


**Supplementary Figure 3. Model simulations for patients with antigen-negative relapse.** Model fits for selected patients that showed relapse of antigen-negative tumor cells. Except for patient G02, the initial tumor burden ($T_N(0)$) was fitted below the detection limit of 2.5$\times 10^5$ cells (gray shaded area).

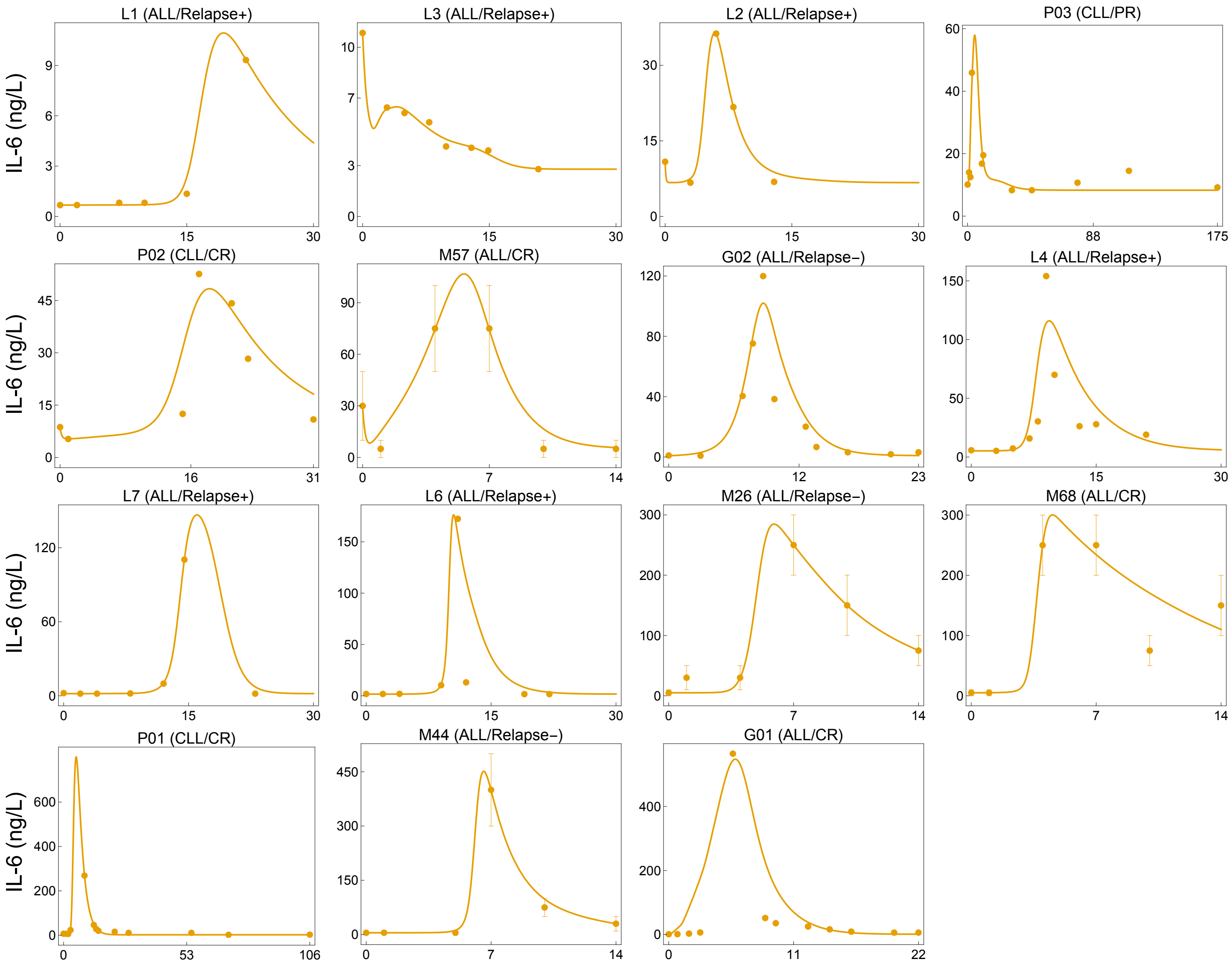


**Supplementary Figure 4. Model simulations of IL-6 kinetics for selected patients**. Experimental data points (orange dots) from [12, 13, 2, 1] are compared with model predictions (orange line). When data was presented data as serum fold change, we establish a direct relationship by considering either a baseline value of 1 ng/L [12] or the specific baseline values provided for each patient [5, 13]. The mean values within the reported range for dataset [1] are visually represented by bars in the figure.

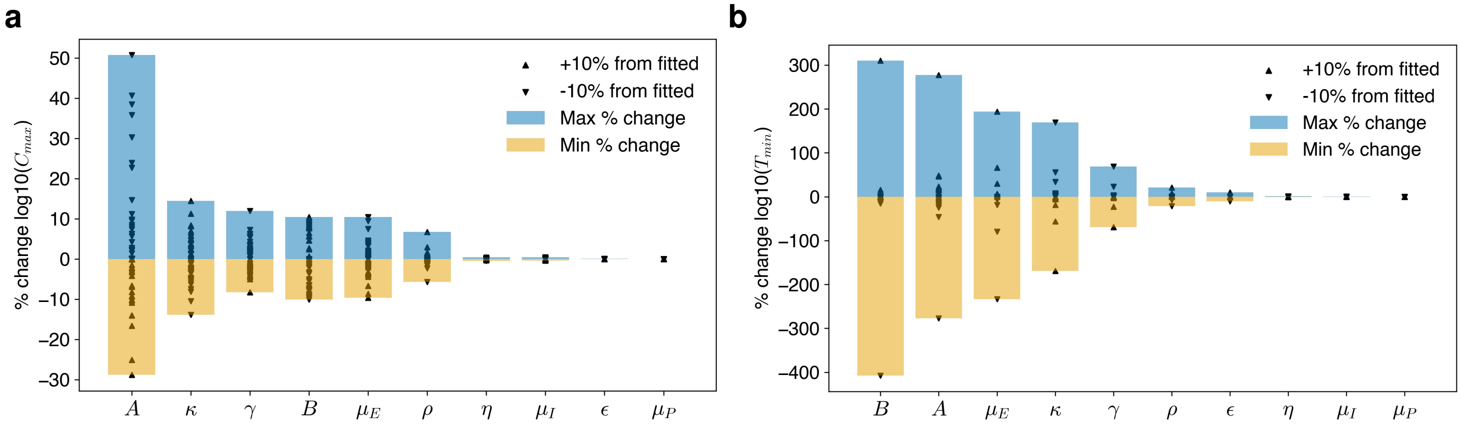


**Supplementary Figure 5. Sensitivity analysis.** The relative change in **a** number of CAR-T cells at peak ($C_{max}$) (all patients) and **b** minimum tumor load (non-responders) achieved during the shrinkage phase ($T_{min}$) were calculated as each parameter varied $\pm 10\%$ at a time. Yellow and blue bars indicate the maximum and minimum percentage changes, respectively. Upward triangles indicate the % change for a 10% increase, while downward triangles indicate the % change for a 10% decrease in each parameter value.

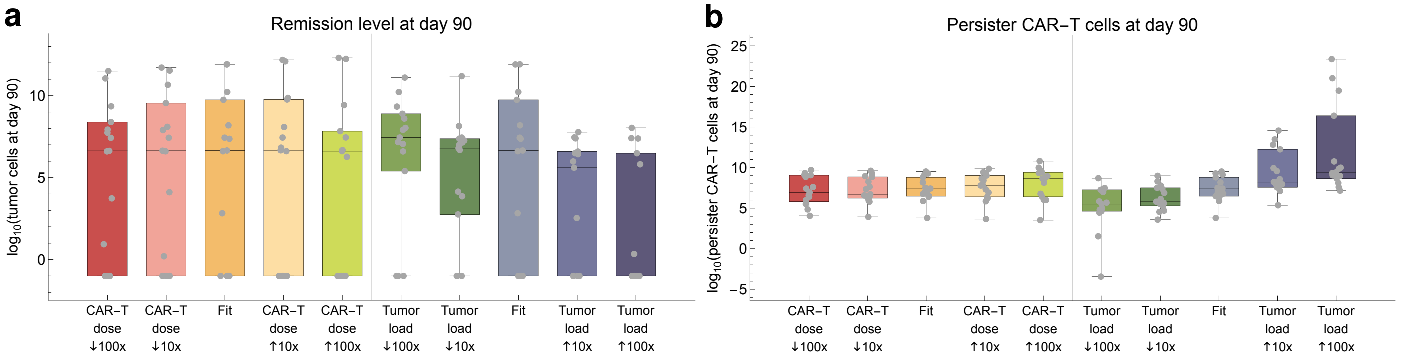


**Supplementary Figure 6. Effect of different dosing protocols on CAR-T cell dynamics and tumor response.** Comparing the standard scenario (Fit) with simulations starting with either a different CAR-T dose or initial tumor burden (10x and 100x smaller and higher). Assessed outcomes: **a** number of tumor cells at day 90, **b** number of persister CAR-T cells at day 90.

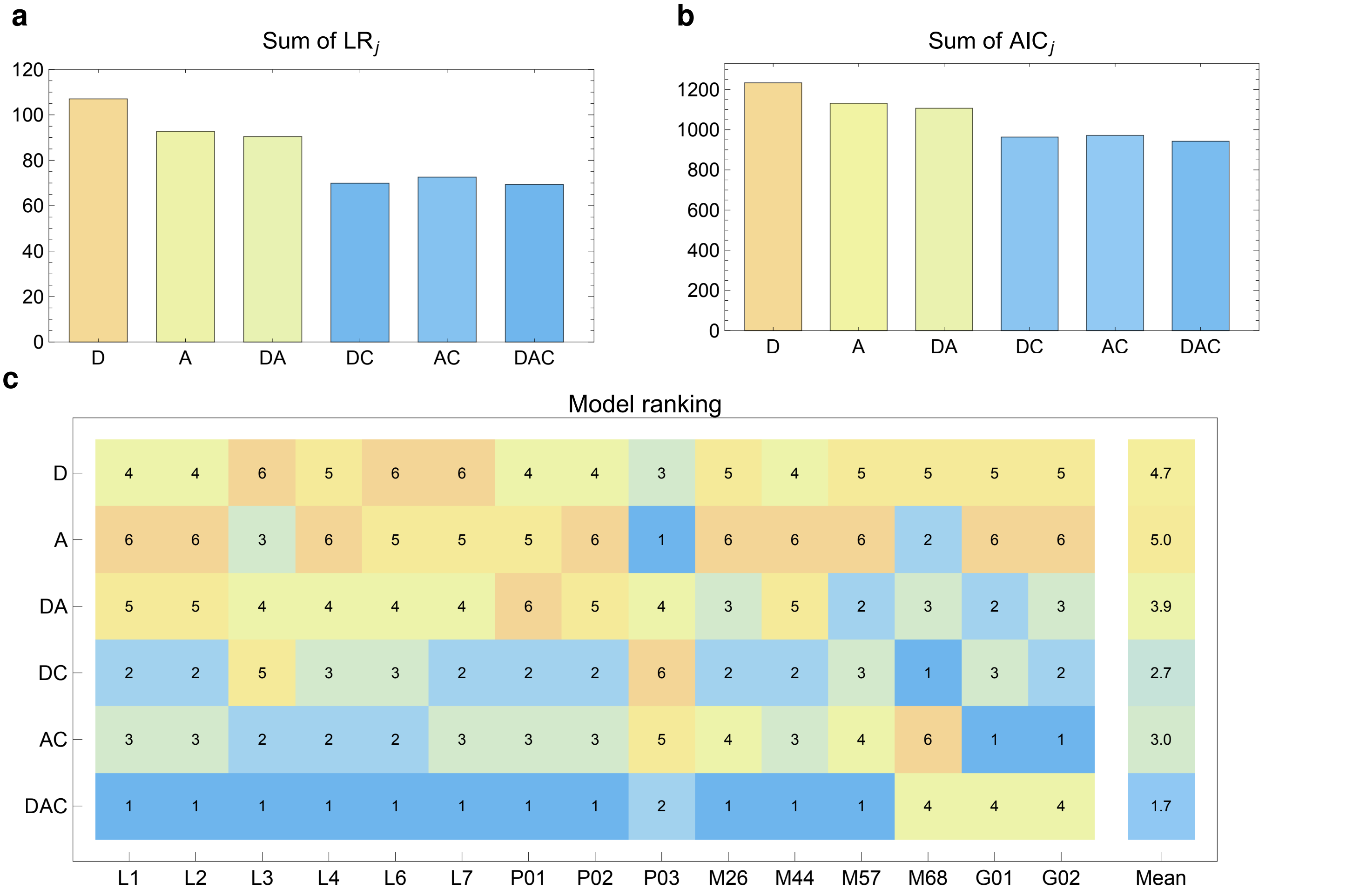


**Supplementary Figure 7. Model comparison.** The automated approach used to estimate macrophage and IL-6 parameters in the model considering three activation mechanisms (denoted DAC - Damps, Antigen, CD40) was applied to the reduced models considering 2 or 1 activation mechanisms (DA, DC, AC, D, A), obtained by setting some $\beta_i = 0$. Model C was not considered because CD40 activation depends on the presence of previously activated macrophages. Panels **a** and **b** show for each model the sum of $LR_j$ (equation (14)) and $AIC_j$ (equation (15)) over all patients. Panel **c** shows the model ranking among patients.

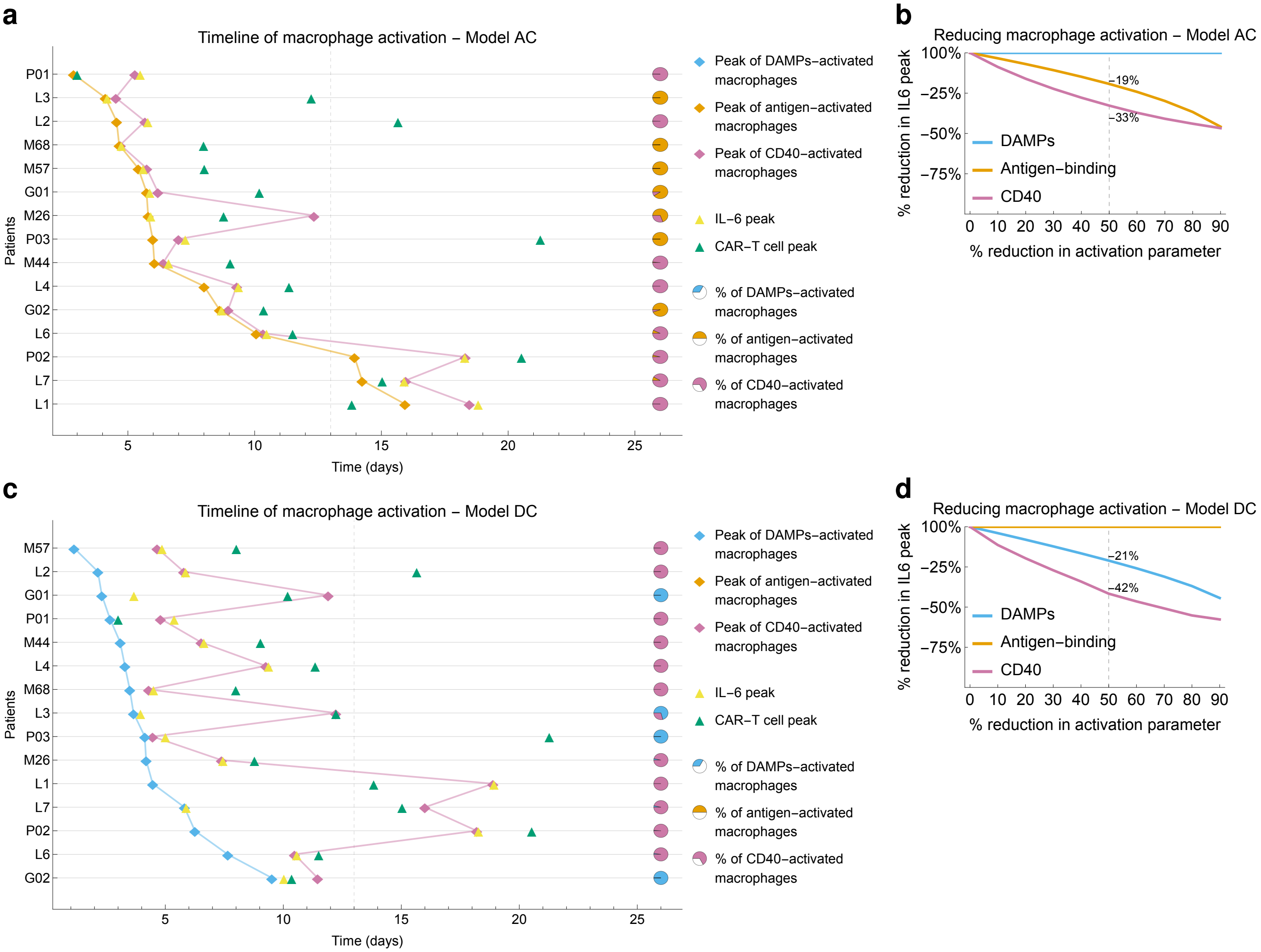


**Supplementary Figure 8. Simulation results for reduced models AC and DC. a,c** Timelines of macrophage-activation, IL-6 and CAR-T cell peaks for each patient. The percentage of CD40-activated macrophages when all patients are combined was 55% and 75% in models AC and DC, respectively. **b,d** Simulation results of interventions reducing mechanisms of macrophage activation one at a time, showing the the overall reduction in IL-6 peak. See Figures 7 and 8 for further details.

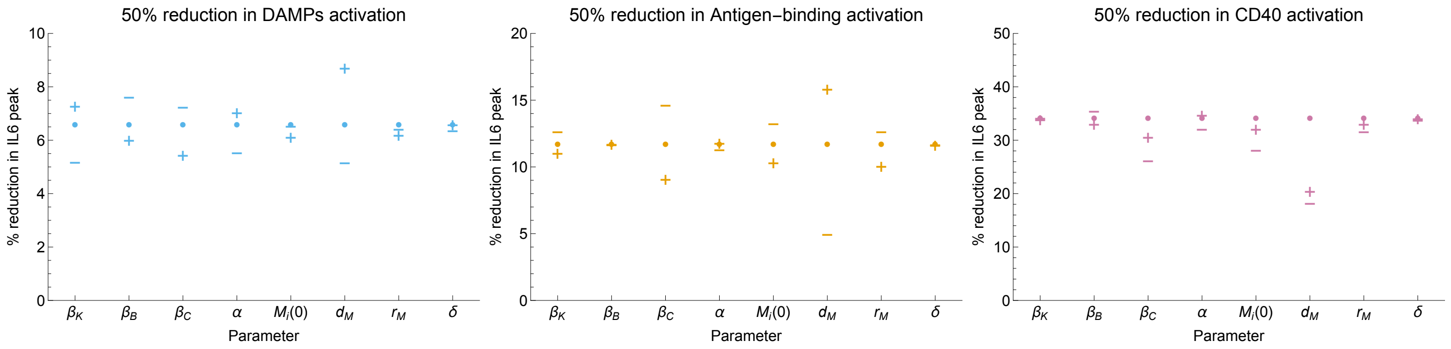


**Supplementary Figure 9. Sensitivity analysis.** A local sensitivity analysis was performed to assess the effect of varying parameters on the % reduction in IL6 peaks. For each patient, a constant 50% reduction in each activation mechanism was simulated and the mean reduction in IL6 peak was calculated (reference scenario, black dots, same values as shown in Fig. 8a fourth panel); this analysis was then repeated by increasing $(+)$ and decreasing $(-)$ each parameter in 50% increments.

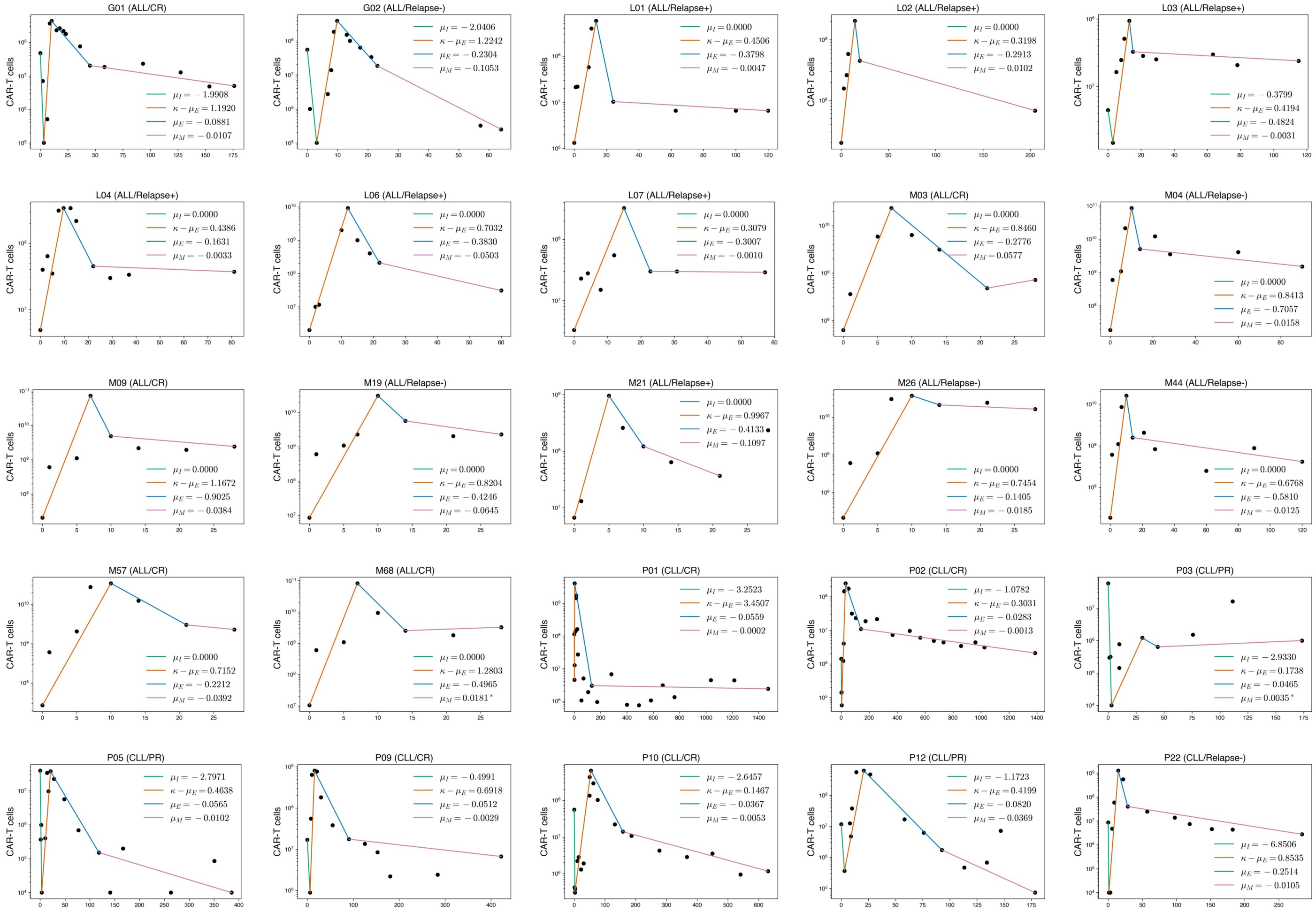


**Supplementary Figure 10. Segmentation of CAR-T cell phases.** Segmentation of CAR-T multiphasic dynamics was performed by defining an exponential curve for each phase. For the distribution phase, we consider data points ranging from the dose until the minimum level of CAR-T cells observed, until 5 days. For the expansion and contraction phases, we consider all data points within the endpoints, which mark the distribution and persistence phases. If the persistence phase is not well marked, we choose the slope that best describes the interior points. Finally, the distribution phase is defined until the last observed data point.

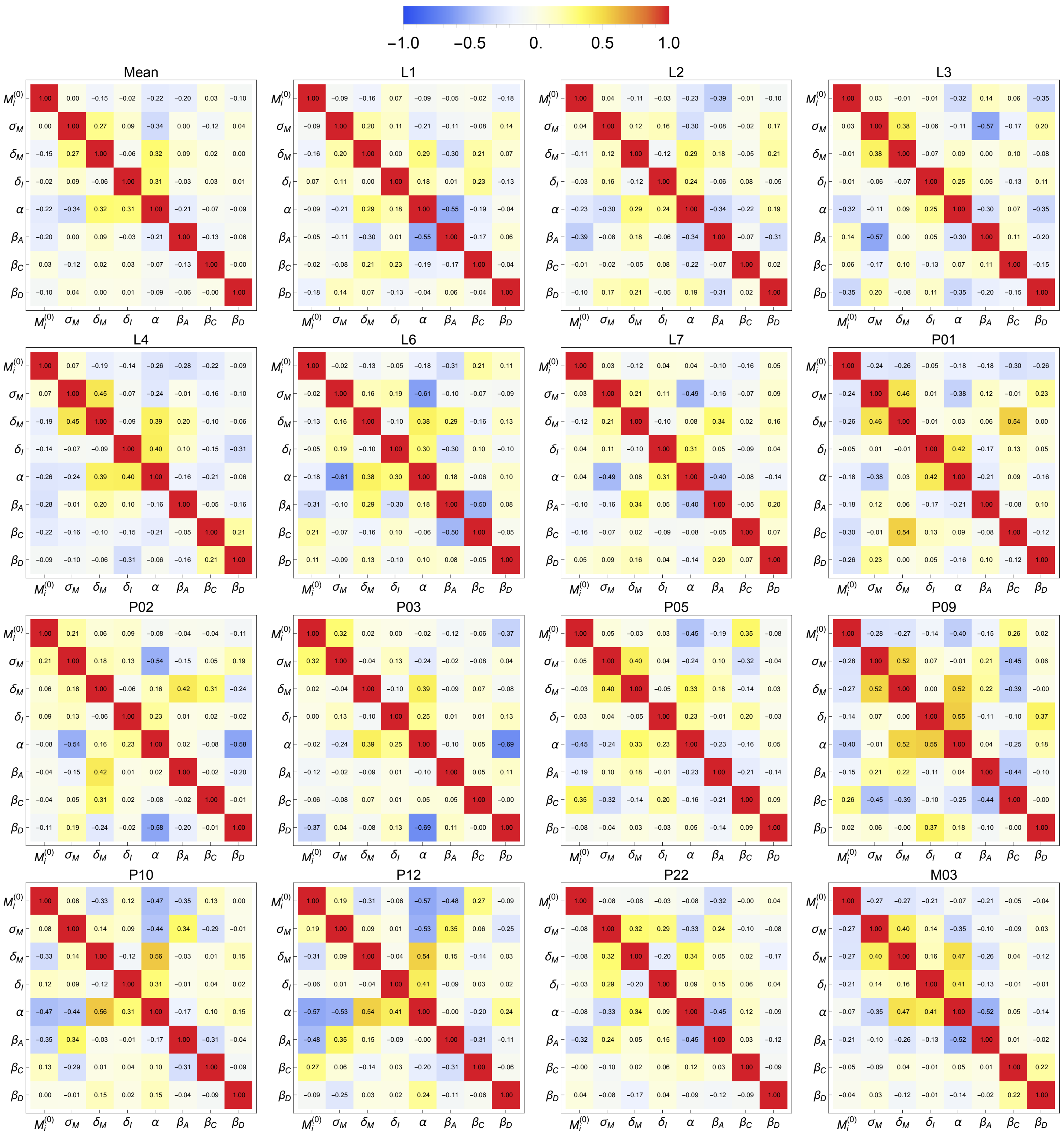

**Supplementary Figure 11. Pairwise correlations for patient-specific IL-6 best fits.** Each plot shows the pairwise correlations between all parameter pairs, considering the 100 best fits obtained after parameter fitting for the IL-6 model, see Supplementary Text S3 for details. The median of all pairwise correlations is also shown.

Supplementary Table 1: Model parameters, their biological meanings, bounds, and references used for parameter estimation.

| Parameter | Unit | Biological meaning | Bounds | Initial guess | REF |
|---|---|---|---|---|---|
| | | | **CAR-T + Antigen Positive** | | |
| $C_I(0)$ | cells | Initial condition for injected CAR-T cells | - | Supplementary Table 2 | Extracted from data |
| $C_E(0)$ | cells | Initial condition for CAR-T expander cells | - | 0 (Fixed) | See "Model Setup" for details |
| $C_P(0)$ | cells | Initial condition for CAR-T persister cells | - | 0 (Fixed) | See "Model Setup" for details |
| $T_P(0)$ | cells | Initial condition for antigen-positive tumor cells | - | Supplementary Table 2 | Extracted from data and refer to "Model Setup" for details |
| $\mu_I$ | day$^{-1}$ | Death rate of injected CAR-T cells | $[0.1-10]\times$slope | $\mu_I = slope$(A) | Extracted from data |
| $\eta$ | day$^{-1}$ | Engraftment rate of injected cells to blood and tumor niche | $10^{-8}-30$ | $\eta = 1$ | [14, 15] |
| $\kappa$ | day$^{-1}$ | Expansion rate of CAR-T expander cells | $[0.1-10]\times$slope | $\kappa = slope$(A) | Extracted from data |
| $\mu_E$ | day$^{-1}$ | Reduction rate (apoptosis and exhaustion) of CAR-T expander cells | $[0.1-10]\times$slope | $\mu_E = slope$(A) | Extracted from data |
| $\epsilon$ | day$^{-1}$ | Rate of long-term memory formation | $2\times[10^{-5}-10^{-1}]$ | $\epsilon = 10^{-2}$ | [7, 16, 14] |
| $\mu_P$ | day$^{-1}$ | Death rate of CAR-T persister cells | $[0.1-10]\times$slope | $\mu_P = slope$(A) | Extracted from data |
| $\rho$ | day$^{-1}$ | Proliferation rate of tumor cells | $[0.0069-0.255]$ | $\rho = 0.1$ | [17, 18] |
| $\gamma$ | day$^{-1}$ | Kill rate of antigen-positive tumor cells by CAR-T expander cells | $0.1-10$ | $\gamma = 1$ | [19, 16, 17](C) |
| $A$ | cell | Half-saturation constant for antigen-receptor binding functional response | $1-4.25\times10^{12}$ | $A = T(0)$ | Assumption based on tumor burden |
| $B$ | cell | Half-saturation constant for tumor kill functional response | $1-1\times10^{10}$ | $B = 10^8$ | Assumption based on maximum CAR-T cell concentration at peak |
| $\theta$ | day$^{-1}$ | Rate of long-term memory recruitment | $[0-0.121]$ | $\theta = 10^{-8}$ (Fixed)(B) | [20, 21, 19] |
| $K$ | cell | Carrying capacity of tumor cells | $4.25\times10^{12}$ | $K = 4.25\times10^{12}$ (Fixed) | Equivalent 100% of blasts |
| | | | **Antigen Negative** | | |
| $T_N(0)$ | cells | Initial condition for antigen-negative tumor cells | $[1-2.5\times10^5]$ | $10^2$ | Assumption undetectable tumor |
| $g_0$ | - | Reduction factor for the kill rate of antigen-negative tumor cells by CAR-T expander cells | $[10^{-3}-10^{-1}]$ | $g_0 = 10^{-2}$ | [7] |
| | | | **IL-6** | | |
| $IL_6(0)$ | (ng/L) | Initial condition for IL-6 concentration | - | Supplementary Table 2 | Extracted from data |
| $M_i(0)$ | cells | Initial condition for IL-6 inactivated macrophages | $[10^8, 10^{10}]$ | Randomly chosen(D) | [8, 9, 10] |
| $M_a(0)$ | cells | Initial condition for IL-6 activated macrophages | - | 0 (fixed) | See "Model Setup" for details |
| $\sigma_M$ | (ng/L)day$^{-1}$ | Basal production of naive macrophages | $[10^8, 10^{10}]$ | Randomly chosen(D) | Estimated from steady state(E) |
| $\delta_M$ | day$^{-1}$ | Macrophages death rate | $[0.1, 1]$ | Randomly chosen(D) | [22, 23] |
| $\beta_K$ | (day.cell)$^{-1}$ | Macrophage activation rate due to DAMP release | $[10^{-15}, 10^{-10}]$ | Randomly chosen(D) | Order of magnitude in equation (11)(F) |
| $\beta_B$ | (day.cell)$^{-1}$ | Macrophage activation rate due to antigen-binding | $[10^{-13}, 10^{-8}]$ | Randomly chosen(D) | Order of magnitude in equation (11)(F) |
| $\beta_C$ | (day.cell)$^{-1}$ | Macrophage activation rate due to CD-40 contact | $[10^{-13}, 10^{-8}]$ | Randomly chosen(D) | Order of magnitude in equation (11)(F) |
| $C$ | cell | Half-saturation constant for macrophage activation rate due to CD-40 contact | $[10^9, 10^{11}]$ | $10^{10}$ (Fixed) | Assumption based on macrophage population |
| $\delta_I$ | day$^{-1}$ | Cytokine natural decay rate | $[1, 24]$ | Randomly chosen(D) | [19, 24, 25] |
| $\sigma_I$ | (ng/L)day$^{-1}$ | Endogenous cytokine production | - | $\sigma_I = \delta_I \min_i IL_{6i}$ (fixed for each patient) | Estimated from steady state(G) |
| $\alpha$ | (ng/L)(day.cell)$^{-1}$ | Cytokine production rate by macrophages | $[10^{-10}, 10^{-5}]$ | Randomly chosen(D) | Order of magnitude in equation (9)(F) |

(A) See Supplementary Figure 10 and Supplementary Text S4 for details.

(B) Fixed to avoid secondary CAR-T cell expansion from just a single dose [26].

(C) These references use the mass action law for tumor cell killing, which is an approximation, for small $C_E/B$, for the saturation function in (4), with a constant $\gamma/B$.

(D) Refer to "Parameter estimation for IL-6 dynamics" for details.

(E) In the absence of activation, the steady state of inactivated macrophages reaches the steady state $M_i^{ss} = \sigma_M/\delta_M$, giving an estimate $\sigma_M = M_i^{ss}\delta_M$; assuming that $M_i^{ss}$ has a range similar to $M_i(0)$ and that $\delta_M \in [0.1, 1]$, we estimate the range for $\sigma_M$ to be $[10^8, 10^{10}]$.

(F) Ranges for the parameters $\alpha, \beta_B, \beta_K, \beta_C$ are difficult to estimate from biological data; these intervals were chosen after initial simulations and considering the orders of magnitude involved in equations (9) and (11).

(G) In absence of IL-6 release by activated macrophages, IL-6 levels reach the steady state $\sigma_I/\delta_I$, which is therefore a lower bound for the minimum IL-6 level during the response phase, thus we set $\sigma_I = \delta_I \min_i IL_{6i}$ for each patient.

Supplementary Table 2: Individual CAR-T cell dose, the end of analysis, disease, outcome, initial tumor burden and baseline IL-6 concentration. In the split-dose regimen, the CAR-T cell dose was given in 3 fractions with 10% administered on day 0, 30% on day 1, and the remaining 60% on day 3.

| **Patient** | **Regimen** | **CAR-T dose** $(\times 10^6)$ | **Weight (kg)** | **Product** | **End of analysis (days)** | **Outcome** | **Initial Tumor Burden** | **IL-6 (0) (ng/L)** |
|---|---|---|---|---|---|---|---|---|
| | | | | **Grupp et al., [12] - ALL** | | | | |
| G01 | Split-dose | 12 (cells/kg) | 40* | CD28 | 330 | $CR$ | NA | 1 ††† |
| G02** | Single-dose | 1.4 (cells/kg) | 40* | CD28 | 64 | $Relapse-$ | 7.7% $(T_N)$<br>92.3% $(T_P)$ | 1 ††† |
| | | | | **Porter et al., [13] - CLL** | | | | |
| P01 | Split-dose | 1130 (cells) | - | 4-1BB | 1590 | $CR$ | $1.7 \times 10^{12}$ (cells) | 7 |
| P02 | Split-dose | 14.2 (cells) | - | 4-1BB | 1560 | $CR$ | $8.8 \times 10^{11}$ (cells) | 8.7 |
| P03 | Split-dose | 586 (cells) | - | 4-1BB | 180 | $PR$ | $2.75 \times 10^{11}$ (cells) | 10.1 |
| P05 | Split-dose | 392 (cells) | - | 4-1BB | 390 | $PR$ | NA | - |
| P09 | Split-dose | 170.0 (cells) | - | 4-1BB | 422 | $CR$ | NA | - |
| P10 | Split-dose | 56.1 (cells) | - | 4-1BB | 630 | $CR$ | NA | - |
| P12 | Split-dose | 118.0 (cells) | - | 4-1BB | 180 | $PR$ | NA | - |
| P22 | Split-dose | 86.4 (cells) | - | 4-1BB | 300 | $PR$ | NA | - |
| | | | | **Li et al., [2] - ALL** | | | | |
| L01 | Split-dose | 13.3 (cells) | - | CD28 | 120 | $Relapse+$ | 0.0% †† | 0.68 †† |
| L02 | Split-dose | 199.6 (cells) | - | CD28 | 240 | $Relapse+$ | 0.57% †† | 10.85 †† |
| L03 | Split-dose | 44.3 (cells) | - | CD28 | 180 | $Relapse+$ | 2.28% †† | 10.85 †† |
| L04 | Split-dose | 48.8 (cells) | - | CD28 | 90 | $Relapse+$ | 0.12% †† | 5.75 †† |
| L06 | Split-dose | 19.9 (cells) | - | CD137 | 60 | $Relapse+$ | 2.28% †† | 1.765 †† |
| L07 | Split-dose | 33 (cells) | - | CD137 | 60 | $Relapse+$ | 0.65% †† | 2.17 †† |
| | | | | **Ma et al., [1] - B-ALL** | | | | |
| M03 | Single-dose | 1.58 (cells/kg) | 40 | 4-1BB | 734† | $CR$ | 0.34% | - |
| M04 | Single-dose | 0.73 (cells/kg) | 26 | 4-1BB | 99† | $Relapse-$ | 0.80% | - |
| M09 | Single-dose | 1.0 (cells/kg) | 20.5 | 4-1BB | 620† | $CR$ | 0% | - |
| M19 | Single-dose | 0.50 (cells/kg) | 17 | 4-1BB | 95† | $Relapse-$ | 0% | - |
| M21 | Single-dose | 0.30 (cells/kg) | 22 | 4-1BB | 30† | $Relapse+$ | 63% | - |
| M26 | Single-dose | 0.50 (cells/kg) | 43 | 4-1BB | 96† | $Relapse-$ | 0.63% | $\leq 10$ |
| M44 | Single-dose | 0.50 (cells/kg) | 37 | 4-1BB | 120† | $Relapse-$ | 0.26% | $\leq 10$ |
| M57 | Single-dose | 0.50 (cells/kg) | 54 | 4-1BB | 280† | $CR$ | 0.07% | 10-50 |
| M68 | Single-dose | 0.50 (cells/kg) | 21 | 4-1BB | 244† | $CR$ | 4% | $\leq 10$ |

* Estimated based on the patient's age.
** Percentage from total of tumor cells, which was not assessed.
† Estimated based on the progression-free survival day.
†† Estimated using WebPlotDigitizer
††† Cytokine concentrations were displayed as factor change from baseline, and for simplicity in fitting and conversion, we assumed IL-6(0)=1 ng/L
NA = Not assessed.

Supplementary Table 3: Calibrated parameter values, in appropriate units (a.u.), used in the simulations for all patients. Parameters whose values were the same for all patients are: $C_T(0) = C_P(0) = M_a(0) = 0$,$K = 4.25 \times 10^{12}$ cells and $\theta = 10^{-8}$ day$^{-1}$.

| | L01 | L02 | L03 | L04 | L06 | L07 | P01 | P02 | P03 | P05 | P09 | P10 | P12 | P22 | M03 | M04 | M09 | M19 | M21 | M26 | M44 | M57 | M68 | G01 | G02 |
|---|---|---|---|---|---|---|---|---|---|---|---|---|---|---|---|---|---|---|---|---|---|---|---|---|---|
| $C_I(0)$ | 1.33E+06 | 2.00E+07 | 4.43E+06 | 4.88E+06 | 1990000 | 3300000 | 1.13E+08 | 1.42E+06 | 5.86E+07 | 3.92E+07 | 1.70E+07 | 5.61E+07 | 1.18E+07 | 8.64E+06 | 6.32E+07 | 1.90E+07 | 2.05E+07 | 8.50E+06 | 6.60E+06 | 2.15E+07 | 1.85E+07 | 2.70E+07 | 1.05E+07 | 4.80E+07 | 5.60E+07 |
| $T_P(0)$ | 1.00E+07 | 3.71E+08 | 9.76E+08 | 7.53E+07 | 1.01E+09 | 306847686.2 | 1.70E+12 | 8.80E+11 | 2.75E+11 | 1.00E+07 | 1.00E+07 | 1.00E+07 | 1.00E+07 | 1.00E+07 | 1.45E+08 | 3.40E+08 | 1.00E+07 | 1.00E+07 | 2.68E+10 | 2.68E+08 | 1.11E+08 | 1.00E+07 | 1.70E+09 | 1.00E+07 | 9.23E+06 |
| $T_N(0)$ | - | - | - | - | - | - | - | - | - | - | - | - | - | 4.76E+00 | - | 6.26E+02 | - | 9.13E+02 | - | 4.24E+03 | 2.34E+03 | - | - | - | 7.70E+05 |
| $IL_6(0)$ | 0.68 | 10.85 | 10.85 | 5.75 | 1.765 | 2.17E+00 | 7.00 | 8.70 | 10.10 | - | - | - | - | - | - | - | - | - | - | 5.38 | 4.89 | 29.79 | 4.86 | 1.02 | 1.03 |
| $\epsilon$ | 0.0120 | 0.0188 | 0.2000 | 0.0200 | 0.0900 | 0.0480 | 0.0001 | 0.0006 | 0.0200 | 0.0005 | 0.0010 | 0.0010 | 0.0073 | 0.0058 | 0.0087 | 0.0147 | 0.0185 | 0.0282 | 0.1026 | 0.0416 | 0.0101 | 0.0085 | 0.0166 | 0.0033 | 0.1000 |
| $\gamma$ | 0.8046 | 0.7063 | 0.5148 | 0.3687 | 1.3367 | 0.4512 | 9.9454 | 0.3654 | 0.6000 | 0.2377 | 1.1852 | 0.2669 | 0.3835 | 0.8827 | 3.2016 | 0.6935 | 4.4002 | 1.0899 | 1.4694 | 1.1871 | 0.6668 | 1.4946 | 4.5905 | 0.9086 | 1.0474 |
| $\kappa$ | 1.8632 | 0.4519 | 2.1875 | 1.7824 | 4.3000 | 3.5898 | 6.3866 | 0.4591 | 2.1000 | 0.6060 | 10.0000 | 0.4331 | 3.5000 | 1.0811 | 1.3637 | 3.6503 | 2.2345 | 2.2198 | 4.3593 | 4.0000 | 3.9881 | 1.4287 | 3.3019 | 1.1801 | 3.1455 |
| $\mu_P$ | 0.0040 | 0.0067 | 0.0019 | 0.0010 | 0.0735 | 0.0038 | 0.0007 | 0.0008 | 0.0002 | 0.0050 | 0.0022 | 0.0047 | 0.0499 | 0.0084 | 0.0001 | 0.0081 | 0.0043 | 0.0075 | 0.0509 | 0.0225 | 0.0046 | 0.0041 | 0.0016 | 0.0083 | 0.1200 |
| $\mu_E$ | 1.4931 | 0.2293 | 1.8000 | 0.8369 | 2.7240 | 1.9996 | 0.1192 | 0.0355 | 0.1000 | 0.1000 | 0.0847 | 0.1073 | 0.2091 | 0.7150 | 0.3713 | 0.5666 | 0.9000 | 0.7757 | 3.1069 | 0.0652 | 0.3697 | 0.2681 | 2.0000 | 0.1074 | 1.4220 |
| $\rho$ | 0.2394 | 0.1293 | 0.0457 | 0.2000 | 0.1993 | 0.2011 | 0.0158 | 0.0128 | 0.0200 | 0.0370 | 0.0101 | 0.0447 | 0.0189 | 0.0507 | 0.0866 | 0.1337 | 0.0308 | 0.1070 | 0.2341 | 0.0906 | 0.0871 | 0.0709 | 0.0979 | 0.0160 | 0.0386 |
| $B$ | 1.75E+03 | 2.01E+07 | 1.39E+07 | 7.00E+02 | 3.24E+07 | 7.45E+03 | 2.14E+09 | 4.50E+06 | 3.00E+06 | 6.00E+05 | 5.56E+08 | 1.32E+04 | 2.40E+08 | 7.15E+07 | 4.15E+07 | 5.68E+06 | 7.13E+08 | 5.88E+04 | 1.26E+08 | 4.79E+07 | 6.12E+05 | 1.83E+07 | 6.38E+08 | 9.41E+03 | 9.57E+07 |
| $A$ | 1.00E+03 | 1.29E+05 | 8.84E+06 | 1.25E+07 | 9.00E+06 | 2.50E+07 | 4.55E+11 | 3.11E+10 | 9.93E+11 | 1.80E+06 | 6.36E+07 | 5.75E+03 | 3.46E+07 | 1.00E+03 | 1.00E+00 | 5.78E+07 | 1.00E+00 | 1.00E+03 | 5.61E+08 | 6.08E+07 | 2.18E+07 | 1.20E+03 | 1.00E+00 | 1.24E+04 | 1.52E+06 |
| $\eta$ | 5.0000 | 1.4803 | 0.8882 | 0.4006 | 9.50E-04 | 3.57E-05 | 0.0010 | 0.5596 | 0.0011 | 0.0083 | 0.0010 | 1.00E-06 | 0.0184 | 2.5969 | 17.1217 | 0.0010 | 1.0000 | 0.1664 | 24.2303 | 0.0010 | 0.0011 | 4.0000 | 8.1093 | 0.0129 | 0.0010 |
| $\mu_I$ | 0.0000 | 0.0000 | 4.7488 | 0.4737 | 1.5016 | 1.0000 | 0.7144 | 1.1517 | 10.0000 | 7.4203 | 0.2481 | 3.0000 | 5.0000 | 18.8636 | 0.0000 | 0.1173 | 0.0000 | 11.2081 | 0.0000 | 21.3448 | 7.4764 | 0.0000 | 0.0000 | 6.0518 | 14.2586 |
| $g_0$ | - | - | - | - | - | - | - | - | - | - | - | - | - | 0.0512 | - | 0.0127 | - | 0.0721 | - | 0.0151 | 0.0160 | - | - | - | 0.0017 |
| $\sigma_I$ | 0.7165 | 80.426 | 4.2076 | 40.477 | 18.452 | 10.749 | 24.557 | 28.684 | 9.0035 | - | - | - | - | - | - | - | - | - | - | 58.697 | 29.901 | 36.129 | 103.72 | 16.449 | 14.204 |
| $\delta_I$ | 1.0599 | 11.9996 | 1.5065 | 7.7423 | 11.404 | 6.1776 | 11.694 | 5.4121 | 1.0848 | - | - | - | - | - | - | - | - | - | - | 11.739 | 5.9801 | 7.2257 | 20.744 | 16.174 | 16.589 |
| $\alpha$ | 4.277E-09 | 1.17E-07 | 7.515E-07 | 1.145-07 | 1.582-06 | 1.413E-07 | 6.760E-06 | 7.783E-08 | 1.816E-08 | - | - | - | - | - | - | - | - | - | - | 1.117E-06 | 1.621E-06 | 6.224E-07 | 1.740E-07 | 6.394E-06 | 2.070E-06 |
| $\delta_M$ | 0.1077 | 0.5184 | 0.9344 | 0.2574 | 0.4304 | 0.7608 | 0.5680 | 0.1601 | 0.5465 | - | - | - | - | - | - | - | - | - | - | 0.4461 | 0.6286 | 0.7011 | 0.1154 | 0.6352 | 0.5432 |
| $\beta_B$ | 1.41E-13 | 8.931E-13 | 1.288E-11 | 5.073E-12 | 1.771E-11 | 2.926E-10 | 6.118E-13 | 3.598E-12 | 2.662E-11 | - | - | - | - | - | - | - | - | - | - | 9.204E-10 | 9.777E-12 | 8.790E-11 | 1.548E-11 | 9.812E-09 | 9.957E-09 |
| $\beta_K$ | 2.100E-12 | 1.303E-13 | 9.037E-12 | 2.882E-14 | 1.686E-13 | 1.093E-12 | 1.441E-15 | 1.142E-15 | 9.698E-11 | - | - | - | - | - | - | - | - | - | - | 2.839E-13 | 7.038E-15 | 2.463E-13 | 1.933E-14 | 2.705E-12 | 3.035E-12 |
| $\beta_M$ | 2.99E-09 | 9.18E-09 | 3.18E-11 | 9.77E-09 | 8.48E-09 | 7.60E-09 | 6.38E-10 | 5.24E-09 | 6.26E-11 | - | - | - | - | - | - | - | - | - | - | 6.45E-11 | 6.92E-09 | 1.38E-11 | 2.90E-12 | 9.93E-09 | 9.99E-09 |
| $\sigma_M$ | 5.30E+09 | 1.11E+08 | 1.34E+09 | 2.00E+08 | 1.02E+08 | 7.99E+09 | 5.44E+08 | 1.49E+08 | 1.19E+08 | - | - | - | - | - | - | - | - | - | - | 8.93E+08 | 1.04E+08 | 1.09E+08 | 1.58E+08 | 1.15E+08 | 1.01E+08 |
| $M_i(0)$ | 3.51E+10 | 9.35E+10 | 1.88E+10 | 9.31E+10 | 9.84E+10 | 1.69E+09 | 5.10E+10 | 6.08E+10 | 8.33E+10 | - | - | - | - | - | - | - | - | - | - | 1.92E+10 | 9.66E+10 | 9.27E+10 | 5.93E+10 | 9.61E+10 | 9.93E+10 |
